# Nanoplastic-mediated non-B DNA mutagenicity

**DOI:** 10.64898/2026.09.08.750059

**Authors:** Nikola Zlatkov, Iris Kufferath, A.K.M Firoj Mahmud, Laurens Mandemaker, John Olsson, Alejandro Medaglia-Mata, Runze Liu, Marion Pollheimer, Vojtech Bystrý, Ulrike Resch, Florian Meirer, Lukas Kenner

**Affiliations:** Department of Molecular Biology, Umeå University, Umeå, Sweden; Department of Pathology, Medical University of Graz, Graz, Austria; Department of Medical Biochemistry and Microbiology, Uppsala University, Uppsala, Sweden; Institute for Sustainable and Circular Chemistry, Department of Chemistry, Utrecht University, Utrecht, The Netherlands; Institute for Risk Assessment Sciences, Department of Population Health Sciences, Faculty of Veterinary Medicine, Utrecht University, Utrecht, The Netherlands; Kvinnokliniken, Inst. NU-sjukvården, Trollhättan, Sweden; Bioinformatics Core Facility, Central European Institute of Technology, Masaryk University, Brno, Czechia; Center of Physiology and Pharmacology, Department of Vascular Biology and Thrombosis Research, Medical University of Vienna, Vienna, Austria; Institute for Genetics, Cologne Excellence Cluster of Cellular Stress Responses in Aging-Associated Diseases (CECAD), Cologne, Germnay; Department of Pathology, Medical University of Vienna, Vienna, Austria; Christian Doppler Laboratory for Applied Metabolomics, Division of Nuclear Medicine, The Department of Biomedical Imaging and Image-Guided Therapy, Medical University of Vienna; Department of Biomedical Imaging and Image Guided Therapy, Division of Nuclear Medicine, Medical University Vienna, Vienna, Austria; Institute of Pathology, Unit of Laboratory Animal Pathology, Department of Biological Sciences and Pathobiology, University of Veterinary Medicine Vienna, Vienna, Austria

## Abstract

Plastic-derived materials have become persistent ecological contaminants since their industrial introduction in the mid-20th century^1,2^. In that context, nanoplastics (NPs) have been discussed as an emerging, ubiquitous plastic-derived pollutant with unique physicochemical properties^3–8^. Their increased surface-to-volume ratio enhances their adsorption, reactivity, and cellular penetration, driving distinct ecotoxicological behaviours when compared to larger plastic particles^9,10^. In this study, we identify direct NP-induced DNA mutagenesis in *Salmonella enterica*. Using a combination of biochemical and biophysical studies, as well as mutagenicity assays, we show that functionalized and non-functionalized NPs induce mutations, depending on the energy state of the bacteria and the NP surface chemistry, by disrupting base-stacking and base-pairing – an intrinsic property of all DNA. Whole-genome analysis revealed that exposure to NPs alters the mutational spectrum and mutation frequency, while circular dichroism spectroscopy demonstrated NP-induced helical flipping from B- to non-B DNA conformations. These motifs preferentially adopt Z-like or A/B-hybrid structures associated with localised mutagenesis through DNA destabilisation. Our combined data reveal that NP-mediated mutagenesis is based on surface chemistry and DNA topology, linking surface chemistry on nanoplastics to genomic instability *in vivo*. The proposed mechanism redefines the current perspective on nanoplastic toxicity shifting it from an indirect stress to direct macromolecular interactions.

This mechanism provides a molecular framework for understanding how NPs could impose mutation bias, environmental selection pressure, and potential genomic risk across biological systems.

## Introduction

Plastics have become deeply embedded in global ecological systems^11–15^. It has been suggested that plastics can fragment over decades into microscopic and nanoscopic debris, and in turn plastics now permeate every environmental compartment^11–15^. Microplastics (MPs) then represent an intermediate degradation stage rather than the end of the plastic life cycle; they are assumed to further break down into nanoplastics (NPs): a class of contaminants whose nanoscale dimensions endow them with novel physicochemical properties distinct from those of bulk polymers or MPs^8,16^.

There is no clear consensus in the literature about what size range defines nanoplastics; some authors use a maximum of 1 micrometer, others use smaller dimensions^17^. According to IUPAC, NP particles of less than 100 nm in diameter are nanoparticles^18^ and thus may act at the boundary between molecular structures and a solid bulk material with a size of more than 100 nm. On the one hand, these NPs behave as dispersed particles in physicochemical processes and conditions, such as Brownian motion and osmolarity, and are part of the dissolved organic carbon of aquatic ecosystems^19^. On the other hand, especially if they precipitate as larger structures, they can behave as a bulk material. Based on their origin, NPs can be classified as primary and secondary^20^. While the primary NPs are manufactured for specific purposes, the formation of secondary NPs is proposed to arise through both abiotic and biotic degradation processes, as well as can also result from human activities^21–26^. Thus far, limited studies suggest that NPs can be generated relatively fast under environmental conditions (e.g., photo- and thermooxidation, hydrolysis and biofragmentation), which can be also accompanied by the incorporation of heteroelements, such as oxygen^19,24,27^. Besides O, other heteroatoms, such as N, can be presumably incorporated into plastics by upcycling, postpolymerization modifications and degradation^28–33^.

Due to their physicochemical properties, the distribution of NPs in nature is significantly more complex and heavily under-researched in comparison to macro- and microplastics. Studies so far imply that on account of their enhanced penetration potential and hydrophobic forces, NPs demonstrate complex long-distance dispersal-dependent persistence in the main components of the Earth system as well as in organs, tissues, and cells of various organisms. NPs have been detected across the Pacific and Atlantic oceans^12–14^, urban atmosphere^11,34^, and lake sediments^15^. A recent study suggested a total nanoplastic concentration of polyethylene terephthalate (PET), polystyrene (PS) and polyvinyl chloride (PVC) nanoplastics of about 18.1 ± 2.1 mg m^−3^ in the entire water column of the Atlantic Ocean^14^. However, reported NP concentrations in the environment and human samples vary hugely owed to the fact that analytical methods for their detection are still being developed and improved^35–37^. For example, it was reported that a human can arguably ingest 2.4 ± 1.3 x 10^5^ nanoplastic particles by drinking just one litre of bottled water^23,38^. Also, while some plastic containers might release 2.11 billion NPs from a square centimeter upon microwaving, depending on the type of plastic, a tea bag may release from 1.20 × 10^9^/mL to 8.18 × 10^6^/mL NPs upon boiling^22,26,39^.

Based on the current state of research, it is assumed that most organisms, and especially humans, experience a lifetime exposure to NPs, with unforeseeable consequences^14,22,23,26^. Yet, PS nanoplastics (PS-NPs), for example, have been shown to be internalised by cells, causing endothelial and epithelial barrier dysfunction, as well as teratogenic, neurological, developmental, and metabolic diseases in both vertebrates and invertebrates^40–50^. While there has been some evidence suggesting that NPs may elicit genotoxicity in mammalian cells, especially in immune cells, linked to oxidative stress, DNA strand breaks and chromosomal aberrations *in vitro*^51–53^, the underlying mechanism of such genotoxicity has remained unresolved. In particular, it is unknown whether specific polymer characteristics, such as chemistries, charge, and size, can directly interact with DNA to induce defined mutational patterns rather than non-specific damage.

Earlier in-vitro studies on DNA-nanoparticle interactions have shown that nanoparticles, such as poly(L-lysine)-coated silica nanoparticles, and polymers, such as poly(ethylene oxide), polylysine and astramol poly(propylene imine) dendrimers, can affect the geometry, rigidity and overall charge of the DNA chains, as well as influence DNA compaction and coil-to-unusual (globular) folding^54–56^. Another study has shown that short and long DNA fragments interact electrostatically with micropolystyrene (200 nm and 2 µm) resulting in the formation of a corona enhancing the immunostimulatory effects of NPs^57^. However, it is currently unknown what effects the particle plastic composition and size have on DNA size and sequence, i.e., whether NPs can induce changes in DNA conformation, and whether they have mutagenic properties.

Here, we address this question directly by investigating whether, aided by their surface chemistry, 25nm diameter PS-NPs induce mutations via direct physical interaction with DNA. Using *Salmonella enterica* auxotrophic mutants as a highly sensitive, quantitative and reproducible genetic biosensor system (Ames test)^58,59–44^ coupled with whole-genome sequencing, high-resolution electron microscopy, chiroptical and infrared spectroscopy, we examined how NP surface chemistry and the physiological state of the bacteria influence mutagenicity and DNA conformation. We show that under native conditions, only aminated PS–NPs, but not unmodified or carboxylated PS-NPs variants induce base substitutions and frameshifts. On the other hand, carboxylated PS-NPs induced only base substitutions in depolarized bacterial cells. Finally, all types of PS-NPs exerted mutagenic activity in DNA methylation-deficient *Escherichia coli* bacteria. The following biophysical analysis demonstrated that direct NP–DNA contact participates in NP-induced mutagenesis, leading to topology-dependent distortion of B-DNA *in vivo.* These findings reveal a direct, charge- and size-mediated mechanism of nanoplastic genotoxicity and establish a molecular link between polymer surface chemistry, DNA topology, and mutation bias, a connection with far-reaching implications for microbial evolution, environmental resilience, and genomic stability across species including humans. Non-B DNA structures such as Z-DNA and cruciforms are enriched at promoters, replication origins, and oncogenic hotspots in the human genome, where they promote mutagenesis and chromosomal instability^60^. Taken together, our results suggest that nanoplastic-induced DNA conformational stress may represent a previously unrecognised route to mutation accumulation linked to, e.g. the risk of cancer, in exposed organisms.

## Results

### Aminated polystyrene nanoplastics (PS–NH₂) induce base substitutions and frameshift mutations in bacteria in a contact-dependent manner under optimal physiology

To assess the mutagenic potential of NPs based on their size and chemistry (**Fig. S1a-e**), we performed an Ames reversion assay using *Salmonella enterica* strains TA1535 and TA1538 which report on base substitutions and frameshift mutations, respectively. Strain TA1535 carries the *hisG46* allele where codon 46 -ctc- is mutated to -ccc- (Leu->Pro), and strain TA1538 carries the *hisD3052* allele, located next to a -cgcgcgcg- hot spot, and carries -1 frameshift deletion in the 298^th^ codon, i.e., -ccgc- mutated to -cgc- (**Fig. 1a**). Upon mutation reversion, both auxotrophic strains regain their ability to synthesise histidine and become prototrophs that form colonies on his-deficient agar media, which indicates that a stable mutation has taken place. To ensure rapid collision kinetics and complete surface saturation, overcoming the natural electrostatic repulsion of the negatively charged *Salmonella* outer membrane^61^, we tested the activity of NPs in a concentration range from 0.01% to 0.1% (m/V). These concentrations force maximum interaction density within controlled incubation timeframes. They also represent a robust, “worst-case scenario” stress or maximum NP exposure model that can mimic natural phenomena that may already be taking place in the plastisphere, biological sinks for NPs, as well as in industrial and urban plastic pollution source point where the concentration of the nanoplastics can exceed well beyond 100mg/L and be modified ^62,63^. From an analytical standpoint, applying a high concentration ensures crossing the detection threshold of signals detected by IR spectroscopy, electron microscopy, and thermogravimetric analysis. PS physicochemical properties—including size, charge, and surface chemistry—were confirmed by circular dichroism, infrared spectroscopy, and transmission electron microscopy (TEM) (**Fig. S1a-e**), and for the initial cell number of bacteria cultured in lysogeny broth (control) prior NP exposure, see **Fig. S1f-g** samples TA1535 and TA1538 and **Materials and Methods**. To test for the mutagenic potential of NPs, TA1535 and TA1538 bacteria were exposed for 1h to suspensions of aminated (PS-NH_2_), carboxylated (PS-COOH) or non-functionalized PS with diameters of 25nm (**Fig. 1b** exposures designated as 0.01% PS, 0.1% PS, 0.01% PS-NH_2_, 0.1% PS-NH_2_ 0.01% PS-COOH, 0.1% PS-COOH and the spontaneous revertants are represented only with the name of the strain).

**Figure 1:**
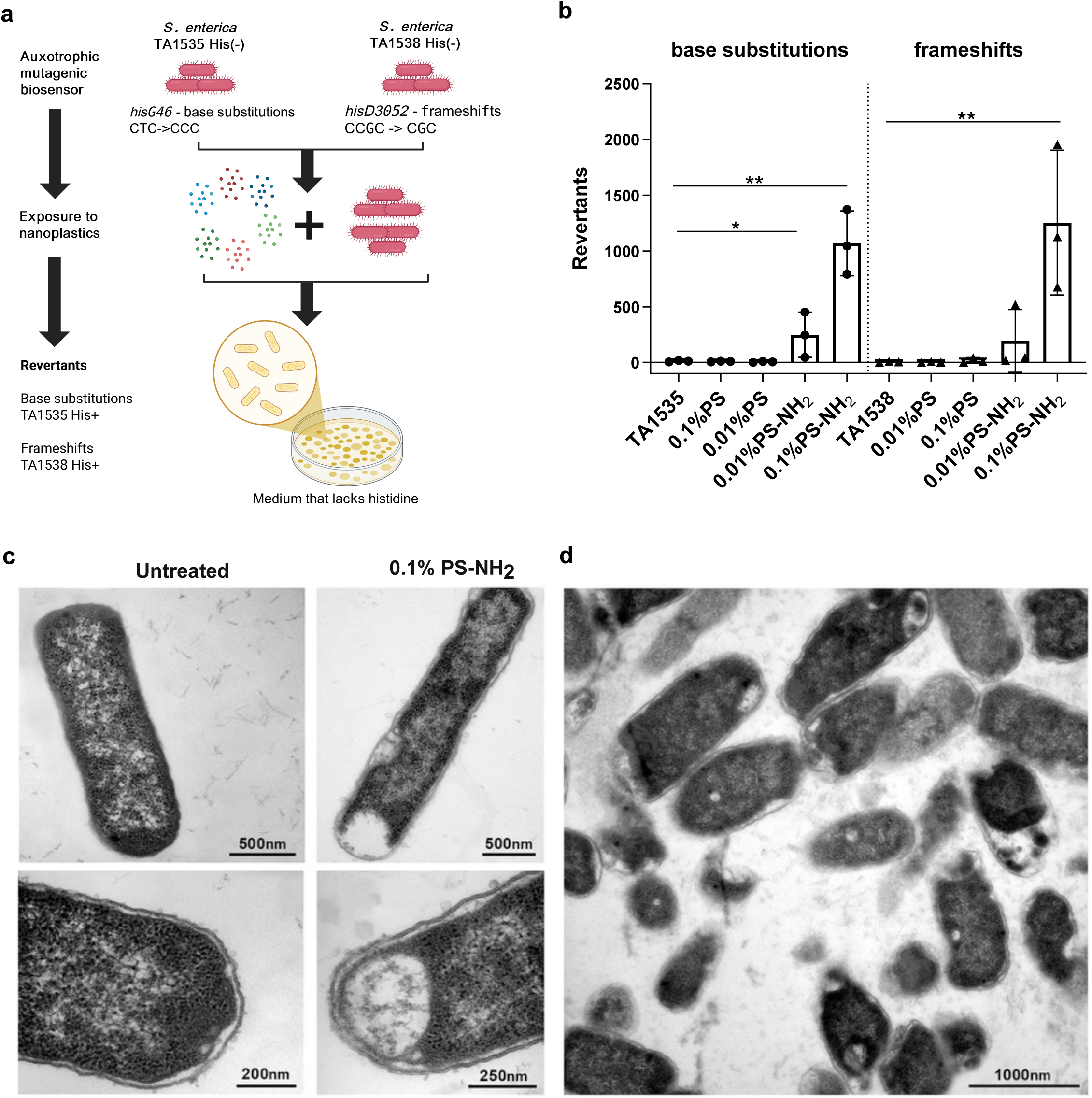
Surface chemistry of polystyrene nanoplastics induces base-substitution and frameshift mutations in *Salmonella enterica ΔuvrB* auxotrophs. **a**, Schematic representation of the Ames reversion assay using *S. enterica* strains TA1535 (base-substitution reporter, *hisG46*) and TA1538 (frameshift reporter, *hisD3052*) to screen for nanoplastic-induced mutagenesis. **b**, Nanoplastic-mediated mutagenesis of strain TA1535 and TA1538 after exposure to 0.01% and 0.1% (w/v) PS, PS–NH₂, PS-COOH. Y-axis indicates the strain and the type of mutation it detects. **c-d**, Transmission electron microscopy (TEM) of *S. enterica* incubated with 0.1 % (w/v) PS–NH₂. **c**, Single cell images of untreated bacteria and bacteria exposed to NPs; **d**, Overview picture of bacteria exposed to NPs.

Remarkably, only the mutants exposed to aminated PS with 25nm diameter generated revertants, i.e., strain TA1535 exhibited 20- and 90-fold increase in revertants (**Fig. 1b** and **Fig. S1h**, see 0.01% PS-NH_2_, P = 0.0372 and 0.1% PS-NH_2_, P = 0.015, unequal variances t-test) while strain TA1538 showed a 250-fold increase only with the 0.1% PS-NH_2_ (**Fig. 1b** and **Fig. S1i** see 0.1% PS-NH_2_, P = 0.011, unequal variances t-test) compared to untreated controls, i.e. the spontaneous revertants. This shows that these positively charged PS–NH₂ NPs promote base substitution at low (0.01%) and high (0.1%) doses, while the frameshift mutations are induced by the PS–NH₂ NPs at a high dose. Moreover, the PS–NH₂ with diameters of 0.3μm (d=0.3 in **Fig. S1h** and **i**) and 1µm (d=1 in **Fig. S1h** and **i**) triggered mutagenic events at their high doses. The presence of 0.1% PS–NH₂ with diameter of 0.3µm increased 10 times the number of revertants of TA1535 (**Fig. S1h**, see 0.1% PS-NH_2_, d = 0.3; P = 0.0091, unequal variances t-test) and 12 times the frameshift revertants (**Fig. S1i**, see 0.1% PS-NH_2_, d = 0.3; P = 0.0283, unequal variances t-test). On the other hand, the presence of 1µm-diameter aminated PS increased only the number of frameshift mutations 10 times (**Fig. S1i**, 0.1% PS-NH_2_, d = 1; P = 0.0173, unequal variances t-test) but not the number of base substitutions (**Fig. S1h**, 0.1% PS-NH_2_, d = 1; P = 0.4737, unequal variances t-test).

So far, the data shows that the mutagenic capability of PS NPs is influenced by their surface properties, i.e., only aminated PS NPs, among those tested, induced base substitution events and frameshifts. The size and dose relationship of aminated PS NPs influence the magnitude and type of the observed mutations. In the case of 25nm diameter NPs, at the lower 0.01% dose, the NPs triggered 20 times higher base substitution events than the baseline control but not frameshifts (**Fig. 1b** and **Fig. S1h-i**). The dose increases from 0.01 to 0.1% also led to a 4-fold increase in the base substitutions. Upon saturation, these NPs induced both base substitutions and frameshifts (**Fig. 1b**). Besides the dose, the size of the NPs also influenced the mutational landscape. Upon a constant mass of 1.5 × 10⁻⁴ g in the samples, the 25nm aminated PS is represented by ∼10^13^ particles, the 300nm aminated PS by 10^10^ particles, and the 1µm aminated PS by 10^8^ particles. This size increase at the expense of particle number led to the reduction in the magnitude of the NP mutagenesis, i.e., the mutagenic effect of the 0.1% 25nm diameter NPs was 8 times higher than that of the 300nm diameter NPs in the case of base substitutions, and 20 times higher than that of the 300nmand 1µm diameter NPs in the case of the frameshift events. Next, to test if the NP backbone chemistry plays a role in the observed mutagenic effects, we exposed the bacteria to 25nm, aminated, non-aromatic poly-(methyl methacrylate) nanoparticles, which led to no mutagenic events (for strain TA1535 see **Fig. S1h** and for strain TA1538 – **Fig. 1i**; samples 0.01% PMMA-NH_2_ and 0.1% PMMA-NH_2_), suggesting that in addition to their surface functionality, dose and size, NP backbone chemistry also play a role in the detected mutagenic effects.

To determine whether the mutagenicity of the NPs depended on metabolic activation to electrophilic species, experiments were repeated in the presence of 2% S9 liver extract. S9 liver extract (the 9000xg supernatant of hepatocyte homogenate) contains cytosolic enzymes and enzymes bound to membrane vesicles derived from the endoplasmic reticulum (microsomes) involved in Phase I and Phase II metabolism of procarcinogens and xenobiotics^64^. When included, S9 supplementation abolished the 25nm PS–NH₂-induced mutagenic response, returning revertant frequencies to baseline (**Fig. S1g**-**h**). This finding hinted that the mutagenic effect might be due to direct PS-NH_2_–bacterium interactions, rather than metabolite-mediated processes. This S9-dependent reduction is consistent with prior evidence that protein corona formation on NP surfaces can shield electrostatic charge and reduce cellular contact^65^, mitigating genotoxicity. Taken together, the reduction of mutagenicity by S9 to basic levels and the selective activity of PS–NH₂ strongly indicate that physical nanoparticle–cell contact, rather than internal metabolism, underlies NP-induced mutagenesis.

Since the inclusion of the S9 microsomal fraction abolished the mutagenic response observed in the Ames test (**Fig. S1j-k**), we aimed to test whether PS-NH_2_ nanoparticles can be taken up by the bacteria. To achieve that, we performed high-resolution transmission TEM. TEM data revealed that the mutagenic PS-NH_2_ NPs were internalised by both strains (**Fig. 1c-d**).

### NPs affect the frequency of spontaneous mutants via mutational shift and structural hotspots

Since only the 25nm diameter PS-NH_2_ NPs induced dose-dependent mutagenic effects, leading to base substitutions (at 0.01 and 0.1% doses) and frameshifts (0.1%), we examined whether PS–NH₂ NPs alter the genomic landscape of spontaneous mutations. To do that, we performed whole-genome sequencing (WGS) on NP-induced revertants (referred to as TA1535+PS-NH_2_ and TA1538+PS-NH_2_), and spontaneous revertants (referred to as TA1535 and TA1538) in ten independent isolates from each group to high coverage (**Fig. 2a**). Across both strains, the overall mutation counts were comparable and profiles were comparable between NP-exposed and control revertants (**Fig. 2b-d**), indicating that PS–NH₂ did not elevate the global mutational burden. Overall, the number of indel events remained low (**Fig. S2a**). On average, the TA1535 spontaneous revertants contained 238 base substitutions and one indel; TA1535+PS-NH_2_ carried 236 base substitutions and 2 indels; TA1538 spontaneous revertants harboured 203 base substitutions and 2 indels, while 1538+PS-NH_2_ showed a reduction to 114 substitutions and one indel (**Fig. 2d** and **Fig. S2a**). While the TA1535 spontaneous and NP-induced revertants showed similar mutation burden, the mutation spectra diverged between the TA1538 spontaneous and NP-induced revertants (**Fig. 2d** and **Fig. S2d**). The PS-NH_2_-induced TA1538 revertants showed a marked reduction in transversion mutations—particularly C->A, G->T, C->G and T->A (**Fig. 2d** and **Fig. S2c-d**). This pattern implies a selective suppression of oxidative-stress-related lesions and suggests that PS–NH₂ exposure favours sequence-directed DNA damage. In contrast, although both strains are derived from *S. enterica* LT2^59^, TA1535PS-NH_2_ revertants kept a substitution spectrum similar to controls (**Fig. S2b** and **Fig. S2c**). To check if NPs trigger specific mutations, we compared the unique mutations discovered only in the NP-induced revertants to the unique mutations discovered in the spontaneous revertants. While the unique mutation number was similar in the NP-induced and spontaneous revertants of TA1535 (**Fig. S2c**), the mutation burden of the NP-induced TA1538 revertants remained lower in terms of transversions (**Fig. S2d**).

**Figure 2:**
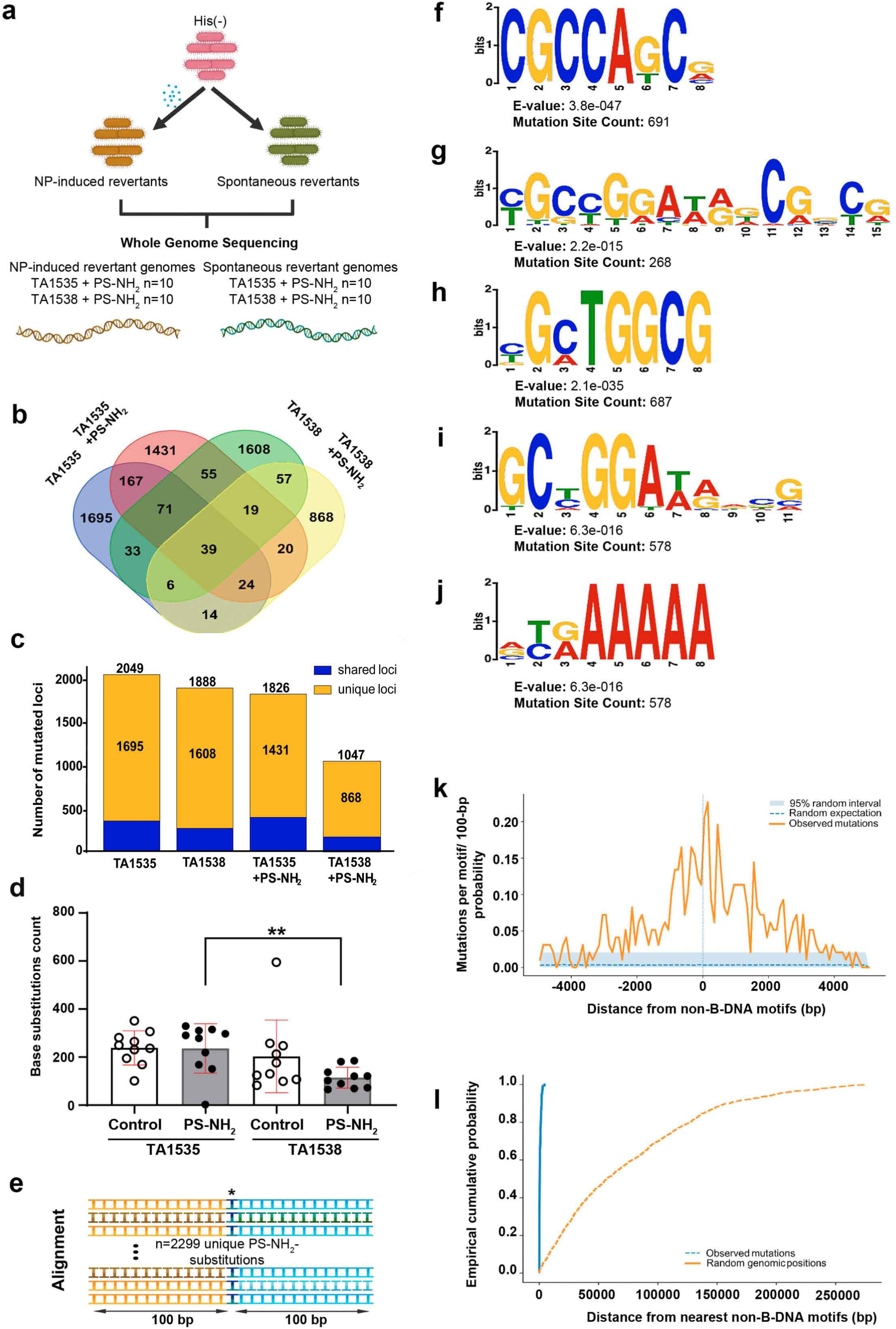
Whole-genome sequencing of spontaneous and NP-induced revertants. **a,** Schematic representation of the revertant comparative genomics. **b**, Venn diagram illustrating the overlap of whole-genome mutation profiles between spontaneous revertants (indicated as TA1535 and TA1538) and nanoplastic-induced revertants (indicated as TA1535 + PS–NH₂, TA1538 + PS–NH₂). **c**, shared (in blue) loci among the spontaneous (TA1535 and TA1538) and NP-induced (TA1535+PS– NH₂ and TA1538+PS–NH₂) revertants, as well as the mutated unique loci (in orange). **d**, Base-substitution spectra of the spontaneous (control) and NP-induced (PS–NH₂) revertants of TA1535 and TA1538. **e**, Schematic representation of the multiple alignment of the upstream and downstream sequences from the unique NP-induced substitutions. **f-g**, Conserved CG-rich motifs 100 nt upstream of the unique mutation sites discovered at the maximum number of mutation sites – n = 691 (**f**) and n = 268 (**g**). **h-i**, conserved CG-rich motif 100 nt downstream of the unique mutation sites discovered at the maximum number of mutation sites – n = 687 (**h**) and n = 578 (**i**). **j**, Conserved polypurine AAAAA motif 100 nt downstream of the unique mutation sites discovered at the maximum number of mutation sites (n=610). **k**, mutational density of the genetic regions with unique NP-induced mutations around non-B-DNA. **l**, empirical cumulative probability of genetic regions with unique NP-induced mutations around non-B-DNA motifs.

We next investigated whether the nanoparticle-induced specific mutations preferentially occurred within specific local DNA sequence contexts (**Fig. 2e**). To do that and to acquire sufficient sequence information for robust estimation of local nucleotide composition, we aligned 100bp upstream (-) and 100bp downstream (+) of the site of unique mutations and checked for a GC bias. The sequence analysis revealed that in the case of TA1538, PS-NH_2_ preferentially induced mutations in the vicinity of DNA sequences with increased GC-content (median GC fraction 0.54 versus 0.53; two-sided Mann–Whitney U test, P = 7.45 × 10⁻³, FDR-adjusted q = 0.0447; **Fig. S2e**-**f**). Moreover, *de novo* motif discovery of the same flanking regions identified highly significant recurrent sequence motifs in both upstream and downstream sequences. The predominant upstream motif was GC-rich (MEME E-value = 3.8 × 10⁻⁴⁷; 691 mutation sites), with a second GC-rich motif also significantly enriched (E-value = 2.2 × 10⁻¹⁵; 268 mutation sites) (**Fig. 2f-g**). Similarly, downstream sequences contained two GC-rich motifs (E-values = 2.1 × 10⁻³⁵ and 6.3 × 10⁻¹⁶; detected in 687 and 578 mutation sites, respectively), together with a highly enriched adenine-rich (poly(A)) motif (E-value = 1.2 × 10⁻²⁵; 610 mutation sites) (**Fig. 2i-j**). Collectively, these findings indicate that nanoparticle-induced mutations are preferentially associated with specific GC-rich and purine-rich sequence environments rather than occurring randomly throughout the genome.

Finally, we analyzed the distribution of 6128 unique locus counts of mutations induced by the presence of NPs and compared them to the unique mutations identified in the spontaneous mutants. Along the chromosome, the periodicity of NP-induced and spontaneous mutations is interrupted by clusters of mutations typical for both groups. Interestingly, the unique NP-induced mutation clusters are in proximity to nucleotide sequences known to trigger non-B structural DNA transitions (**Table S1**). Genomic coordinates of predicted non-B-DNA-forming motifs and single-nucleotide substitutions were used to find out whether mutations occurred preferentially in the close proximity of non-B-DNA sequence elements. Observed mutation sites were significantly closer to predicted non-B-DNA motifs than randomly sampled genomic positions (**Fig. 2k**, median distance, 386 bp versus 57.2 kb; one-sided Mann–Whitney U test, *P* = 2.79 × 10⁻⁸⁹). This indicates a strong, non-random positional association between mutations and non-B-DNA-forming sequence elements, also indicated by their cumulative probability (**Fig. 2l**). Taken together, the fact that NPs increase the frequency of revertants without affecting the mutational burden and that NPs induce mutational shift leading to the formation of mutational clusters in proximity to non-B-DNA elements indicates that NPs trigger mutagenesis via a topology-based mechanism.

### Polystyrene nanoplastics physically associate with chromosomal DNA

Unlike eukaryotic chromosomes, the prokaryotic genetic material is not separated from the cytoplasm by a nuclear membrane, and it is localized directly in the cytoplasm as a condensed nucleoid. Therefore, to explain the NP-induced changes in bacterial genetics, we tested if NPs uptaken by the bacteria (**Fig. 1c-d**) can interact directly with their chromosomes. To achieve that, in combination with TEM, infrared spectroscopy (IR) and thermogravimetric analyses, we performed bacterial cell fractionation in which we first isolated the bacterial periplasmic fraction from the bacterial protoplasts and from the bacterial protoplasts we precipitated the fraction that contained the NP-DNA complexes (see **Fig. 3a** for experimental design).

**Figure 3:**
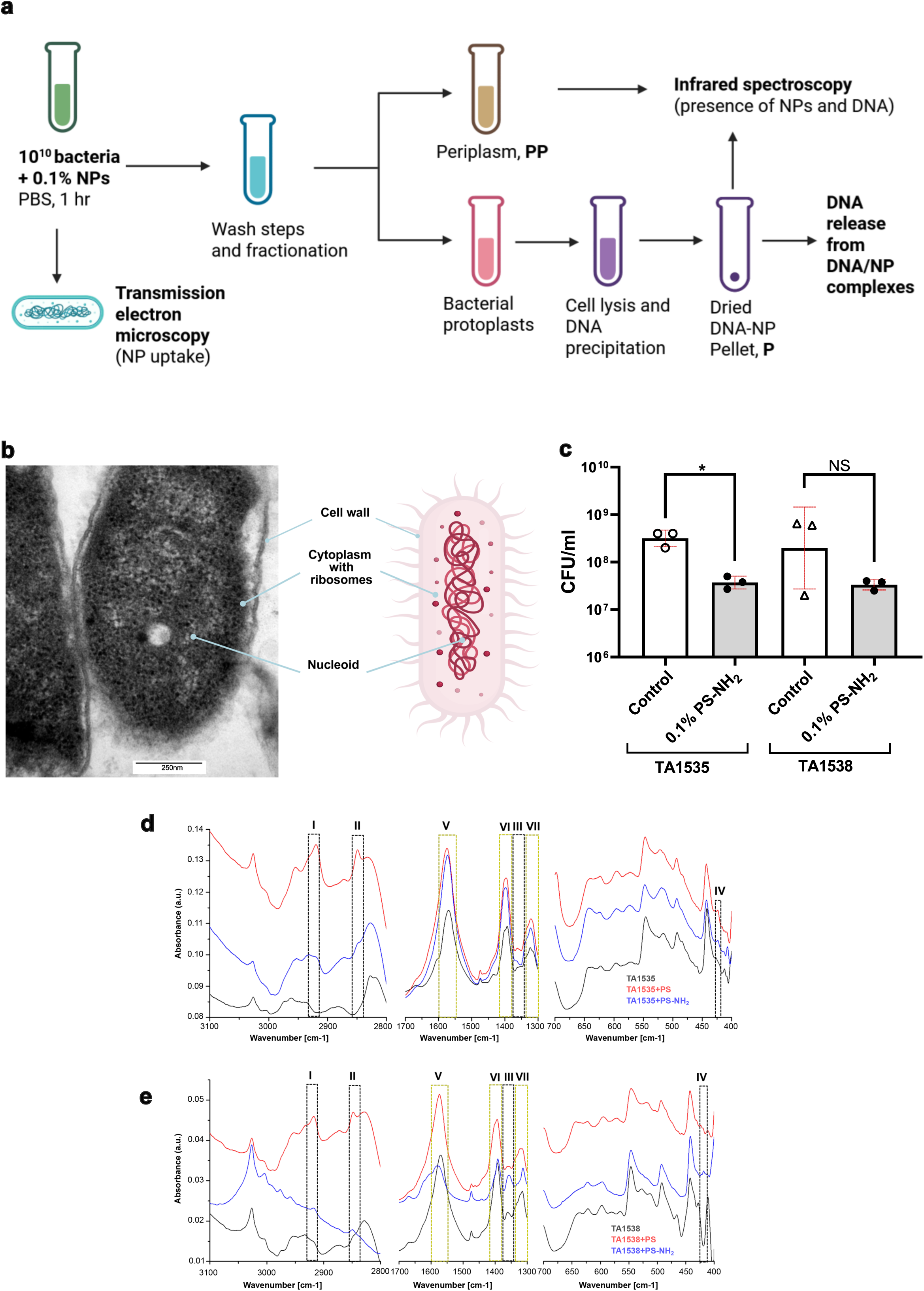
Internalized polystyrene nanoplastics within *S. enterica* bacteria associate with chromosomal DNA. **a,** Schematic overview of the experimental setup of NP detection in bacteria (PP-periplasm, P – pellet enriched with bacterial DNA). **b**, colocalization of NPs with the bacterial chromosome. **c**, Viable cell counts (CFU ml⁻¹) of bacteria incubated with and without PS–NH₂. **d-e**, Infrared spectroscopic analysis of cytosolic DNA fractions from strain TA1535 (**d**) and TA1538 (**e**) exposed to PS and PS–NH₂ compared with unexposed controls. Spectra display characteristic aliphatic C–H stretching (boxes I, II), C–C stretch and skeletal vibrations of the PS backbone (boxes III, IV), alongside canonical DNA phosphate and base–sugar modes (boxes V, VI and VII; ∼1238 cm⁻¹ asymmetric, ∼1080 cm⁻¹ symmetric PO₂⁻ stretching, ∼ 1300 cm⁻¹ base-sugar vibrations). In black, unexposed control DNA; in red, PS-exposed; in blue, PS–NH₂-exposed samples; x axis – wavelength, cm^-1^ and y axis - absorbance).

Interestingly, the TEM experiments showed that several particles appeared to co-localise with condensed nucleoid regions, suggestive of direct polymer–DNA contact (**Fig. 1c-d** and **Fig. 3b**). Viability assays showed no significant reduction in CFU/ml between exposed and control cultures, suggesting that the NP-carrying bacteria are alive (**Fig. 3c**, compare TA1535 to TA1535 0.1% PS-NH_2_ and TA1538 to TA1538 0.1% PS-NH_2_), confirming that NP exposure caused genotoxic but not cytotoxic effects when bacteria are in their optimal physiology state. To a lesser extent, bacteria were able to take in also PS NPs and PS-COOH NPs but the NPs were associated predominantly with the outer surface of the outer membrane (**Fig. S3a** boxes I, II and V and **Fig. S3b** box II) and the NPs which were taken in, showed the tendency to be located away from the bacterial chromosome within the bacterial cytoplasm (**Fig. S3a** box III, **Fig. S3b** box I, II, III, IV and V).

To verify whether the NPs co-localise with the bacterial chromosome, we isolated subcellular fractions, i.e., extracellular milieu, periplasmic contents, and cytoplasmic fractions enriched in genomic DNA, and analysed them by infrared spectroscopy (**Fig. 3d-e** and **Fig. S3c-d**). PS and PS-NH_2_ characteristic vibrational peaks (asymmetric and symmetric C–H stretch at ∼2915 cm⁻¹ and 2848 cm⁻¹, C–C stretch near 1450 cm⁻¹, and skeletal ring vibrations around 700–750 cm⁻¹) were detected as expected in the extracellular fraction of exposed bacteria (**Fig. S3c**). Signal intensity in the periplasmic fraction was negligible (**Fig. S3d**), indicating that most surface-bound or not internalised NPs were effectively removed during washing and fractionation.

The fraction with bacterial chromosomes showed IR vibrations typical for PS in both TA1535 and TA1538 bacteria (**Fig. 3d-e**, boxes I, II, III, IV). The genomic DNA fraction exhibited distinct spectral signatures corresponding to the canonical control peaks (PO₂⁻ asymmetric stretch at 1238 cm⁻¹; symmetric stretch at 1080 cm⁻¹; **Fig. 3d-e**, boxes V, VI and VII; see **Fig. S3e** for control DNA measurement). The presence of aromatic and aliphatic C–H vibrational bands unique to PS confirmed co-isolation of NPs with the chromosomal DNA fraction, consistent with NP–DNA association rather than extracellular contamination. Notably, PS–NH₂ spectra displayed stronger PS-specific absorbance and broader DNA-associated peaks compared with non-functionalized PS, implying enhanced electrostatic or hydrogen-bond-mediated binding between the cationic PS–NH₂ surface and negatively charged DNA phosphate backbones (**Fig. 3d-e**). In both strains, PS- and PS–NH₂-exposed samples show increased intensity in polymer-associated spectral regions while retaining DNA-specific signatures. The consistent detection of polymer-derived vibrational features within DNA fractions supports co-isolation and physical association of PS-NPs with genomic material, rather than direct chemical modification or disruption of DNA structure.

To further validate that the aminated PS NPs interact significantly more with the bacterial genome than the pristine PS NPs, we performed DNA release assay in which we tested the extent to which the bound DNA in the DNA-NP complexes dissolves in water in comparison to the DNA pellet extracted from the untreated bacteria. As shown in **Fig. S3f**, the amount of DNA released from the pellet highly depends on the incubation of the bacteria with NPs and the type of NPs. The highest amount of DNA was released by the control sample (average 261.833ng/µl), followed by the DNA released from the DNA-PS complex (143.5 ng/µl) while the least amount of DNA was released from the DNA-PS-NH_2_ complex (63.783ng/µl). This experiment demonstrated that: first, there are intracellular interactions between the bacterial chromosome and the uptaken NPs, and second, the types of interactions are defined by the surface chemistry of the NPs.

Taken together, these results demonstrate that nanoplastics—particularly positively charged, aminated PS-NPs—can penetrate viable bacterial cells, localise near the nucleoid, and physically associate with genomic DNA, providing a plausible mechanism for the base-substitution and frameshift mutagenesis observed in **Fig. 1**. Our data also confirm and extend previous findings that nanoplastics can be bioaccumulated in bacteria^44,66–68^. Yet, our data further indicate that surface functionalization and NP size critically modulate nanoparticle-DNA affinity, with positively charged aminated PS showing markedly higher DNA-binding potential than unmodified, uncharged PS. These findings support a novel mechanism of NP-induced genotoxicity based on direct interaction with the bacterial genome.

### Collapse of membrane potential increases the uptake of PS nanoplastics and promotes base substitution mutations induced by carboxylated PS nanoparticles

To probe whether the observed NP internalisation and NP mutagenicity are dependent on bacterial energetic defences, *S. enterica* TA1535 cells were pre-incubated with 50µM carbonyl cyanide m-chlorophenyl hydrazone (CCCP). CCCP is a protonophore that collapses the proton motive force and decouples the transmembrane electrochemical gradient, thereby placing the bacteria into a low-energy state with disrupted membrane potential, which also leads to protein mislocalisation^69^. After CCCP pre-treatment, bacteria were exposed to 0.1% (w/v) PS-NPs suspension of aminated (PS–NH₂), carboxylated (PS–COOH), or non-functionalized PS of 25nm diameter, see **Fig. 4a** for experimental setup overview. Remarkably, TEM imaging revealed enhanced NP accumulation under these low-energy conditions (**Fig. 4b-d**), supporting a proton gradient-independent uptake mechanism. In CCCP-treated cells, non-functionalized PS showed peripheral accumulation (**Fig. 4b**, box VI) while PS– NH₂ exhibited three distinct accumulation patterns: (i) complete cytoplasmic plastization (**Fig. 4c**, box I), (ii) dense intracellular NP inclusions (**Fig. 4c**, box II), and (iii) discrete particle localisation near the cell periphery or nucleoid (**Fig. 4c**, box III). PS-COOH-NP exposure led to intracellular localisation similarly to (ii) or (iii) (**Fig. 4d**, box IV and V). These findings demonstrate that together with the membrane potential, surface charge governs NP entry efficiency.

**Figure 4:**
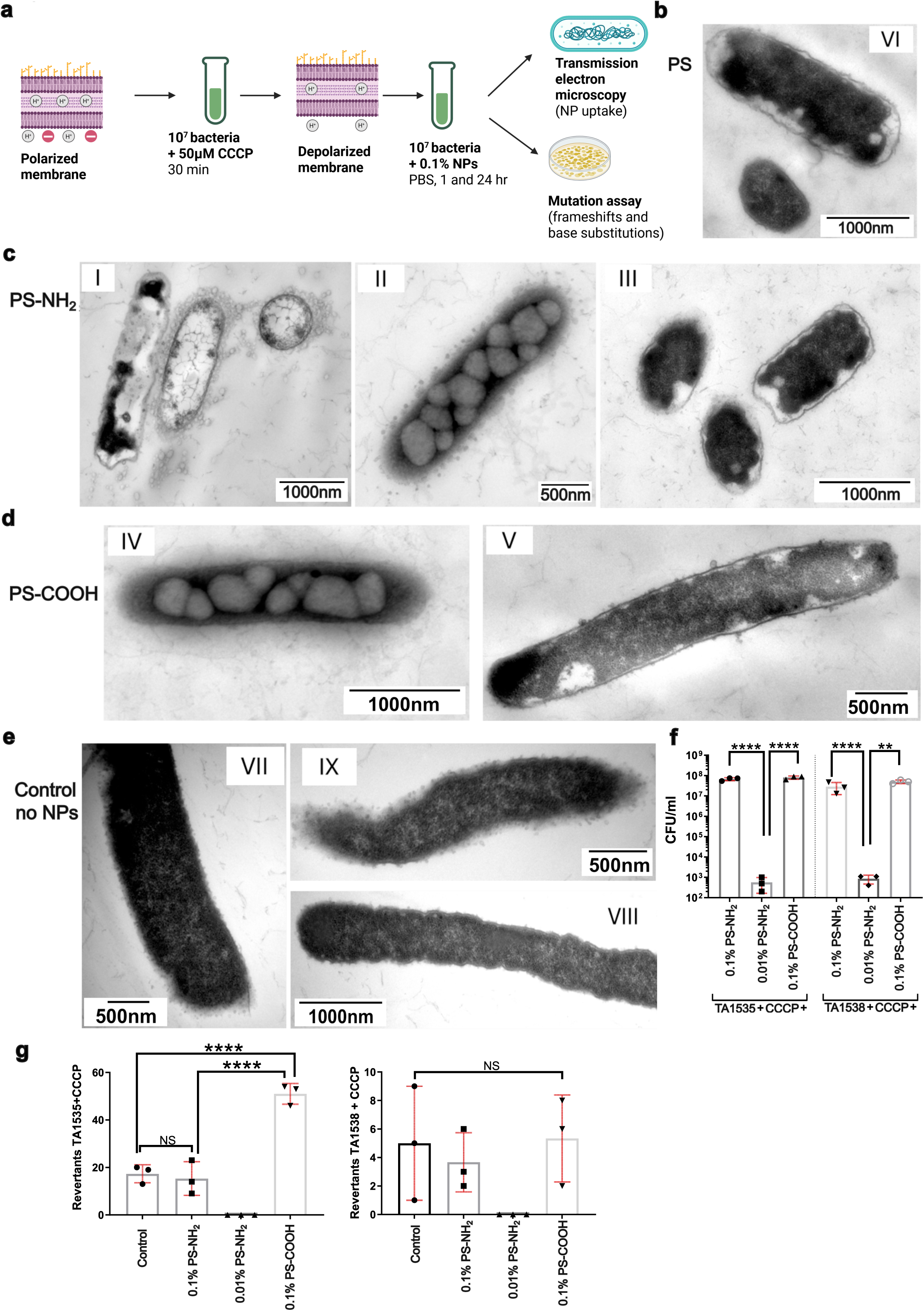
The role of membrane potential in polystyrene nanoplastics (PS-NPs) uptake and their genotoxic outcomes in *Salmonella enterica* TA1535 and TA1538. **a**, Schematic overview of the experimental setup of depolarized bacterial strains exposed to NPs. **b**-**e**, Transmission electron microscopy of *S. enterica* TA1535 and TA1538 pre-treated with 50 µM carbonyl cyanide m-chlorophenyl hydrazone (CCCP) to dissipate the proton motive force, followed by (**b**) 1 h exposure to 0.1% (w/v) non-functionalized PS, (**c**) aminated PS (PS–NH₂), or (**d**) carboxylated PS (PS–COOH). (**e**) Untreated CCCP controls are shown. Under energy-depleted conditions, PS–NH₂ nanoparticles exhibited extensive intracellular accumulation, forming dense inclusions and localized aggregates (boxes I–III), whereas PS–COOH and neutral PS displayed moderate internalization (boxes IV–VI). CCCP-treated, NP-unexposed cells showed slight elongation but intact membranes. **f**, Viable cell counts (CFU ml⁻¹) of CCCP-treated bacteria exposed to 0.1% PS (TA1535+CCCP 0.1% PS and TA1538+CCCP 0.1% PS), 0.1% PS–NH₂ (TA1535+CCCP 0.1% PS–NH₂ and TA1538+CCCP 0.1% PS–NH₂) and 0.1% PS-COOH (TA1535+CCCP 0.1% PS-COOH and TA1538+CCCP 0.1% PS-COOH) after 1h of exposure indicate reduced viability and cytotoxicity under low-energy conditions. **g**, Ames reversion assay of CCCP-treated cells exposed to 0.1% PS, 0.1% PS–NH₂ and 0.1% PS-COOH NPs overnight.

The decoupling of membrane potential appeared to facilitate PS uptake in a fashion similar to that seen under normal physiological conditions, especially for PS-NH_2_ by bacteria with normal membrane potential (compare **Fig. 1c-d** to **Fig. 4b-c**). By contrast, CCCP-treated, NP-unexposed bacteria (**Fig. 4e**) showed slight elongation and loss of turgor as expected^69^, and typical of energy depletion, but no signs of NP-induced morphological disruption confirming that the observed ultrastructural changes in other panels were NP-dependent. Exposure to PS-NH_2_ under these low-energy conditions significantly reduced bacterial viability, amplifying NP cytotoxicity (**Fig. 4f**). Neither PS–COOH nor non-functionalized PS caused significant lethality. To assess whether this altered energy state affected NP-induced mutagenesis, Ames test was performed on CCCP-treated bacteria. Again, the exposure corresponded to an approximate NP-to-cell ratio of 1000:1, respectively (for cell number of the CCCP-treated bacteria prior NP exposure see **Fig. S1f-g** samples TA1535+CCCP and TA1538+CCCP). After one hour of exposure, none of the NPs induced detectable reversion events (**Fig. 4Sa-b**). However, overnight exposure revealed striking differences: PS–NH₂ was lethal, leaving no surviving revertants, whereas PS–COOH increased the revertants by ∼2.5-fold relative to untreated controls (**Fig. 4g**). This unexpected finding suggests that while PS–NH₂ triggers acute cytotoxicity under depolarized conditions, PS–COOH, despite its anionic charge, can induce delayed mutagenic responses when cellular metabolism is suppressed.

To test if the observed effects are due to the surface chemistry of the NPs, we incubated the CCCP-treated bacteria with 25nm PMMA, PMMA-NH_2_ and PMMA- COOH for 24h. The results showed a trend similar to the effect of the PS, PS-NH_2_ and PS-COOH NPs on the CCCP-treated bacteria. PMMA-NH_2_ caused a 1000-fold decrease in the bacterial number in strain TA1535 (**Fig. S4c**) and a 10-fold increase in strain TA1538 (**Fig. S4d**) after 24h. When tested for base substitution and frameshift events, only the CCCP-treated bacteria of strain TA1535 generated revertants after exposure to PMMA-COOH NPs (**Fig. S4d-e**), which is in line with the fact that only the CCCP-treated TA1535 bacteria exposed to PS-COOH generated revertants, indicating that carboxyl surface chemistry affects base substitutions in bacteria in a low energy state.

Overall, these data reveal that the bacteria’s energetic state modulates both the route and consequence of NP entry: under optimal physiological conditions, uptake of PS– NH₂ is efficient and mutagenic but largely non-lethal. When the proton gradient is dissipated, NP uptake is increased, which allows the PS-COOH NPs to interact directly against the bacterial chromosome, resulting in nucleotide alteration.

### NPs distort B-DNA topology

Thus far, our data suggest that NP exposure does not increase the number of mutations but only shifts the mutational burden (**Fig. 1b** and **Fig. 2b-d**), and the taken-up PS-NPs co-localise with the bacterial chromosome (**Fig. 3d-e**). This, together with the identified conserved upstream and downstream elements of the unique mutations (**Fig. 2f-j** and **Table S1**), raised the question of whether the NP-mutagenesis is triggered via DNA interactions that alter the physical architecture of DNA and hence the bacterial genome. Such interactions could distort canonical B-DNA into alternative conformers and are known to trigger mutagenesis. Some of them are the left-handed Z-DNA, the non-canonical triple helical H-DNA the structures of which are shown to be maintained by non-Watson-Crick interactions^70–74^. To test if and what non-B conformations can be induced by NPs, we conducted in-vitro assays exposing linear synthetic heteroduplexes, as well as natural circular DNA, such as bacterial chromosomes and plasmids, to NPs of varying surface chemistries (**Fig. 5a** for experimental design). It should be noted that the spectra of the tested NPs (**Fig. S1a**) show that surface functionalization does not induce detectable chiroptical properties or structural alterations and exclude major aggregation-related artefacts.

**Figure 5:**
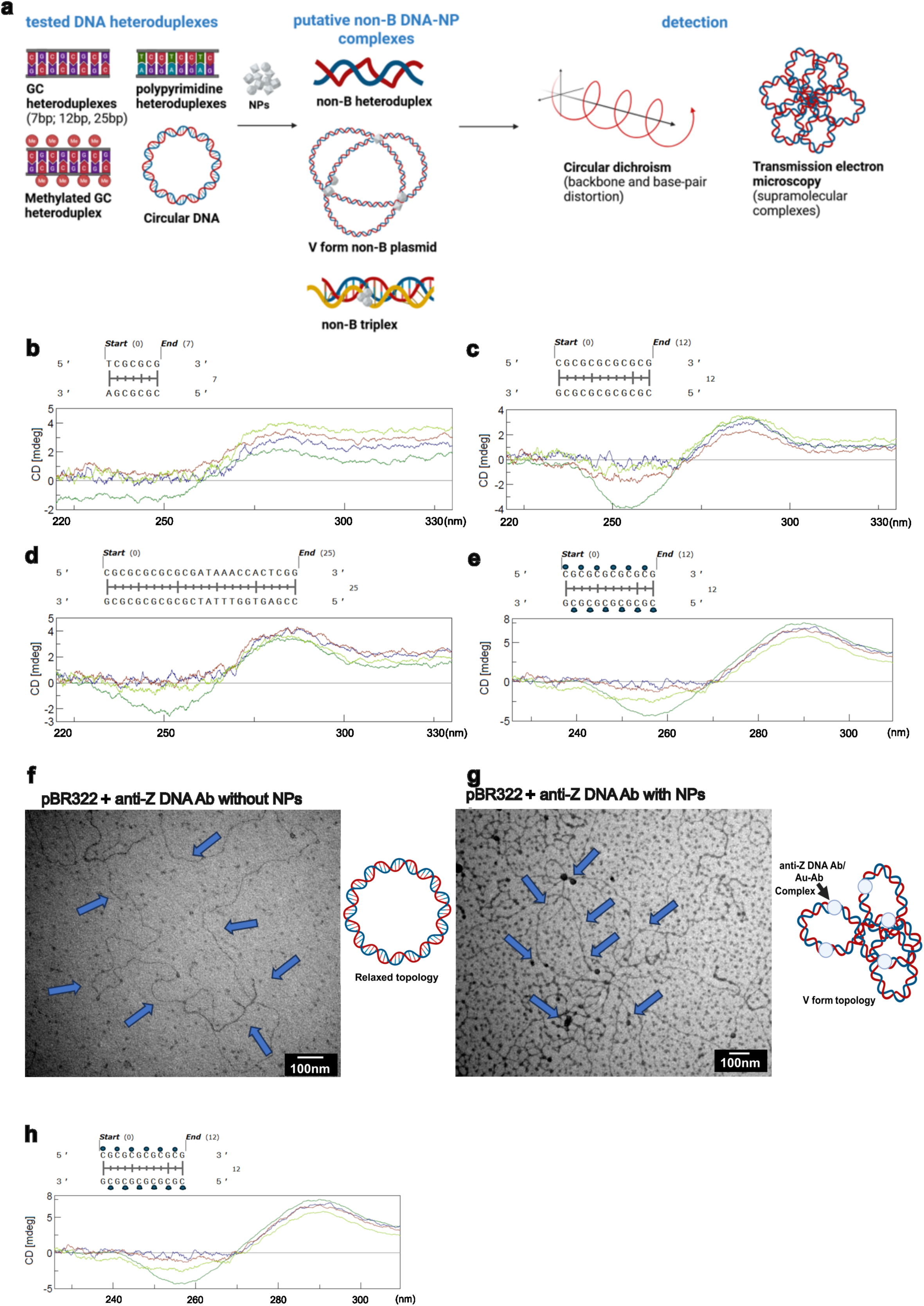
The role of DNA sequence, length, and shape in nanoplastic-DNA interaction. **a**, Schematic overview of the tested DNA sequences for interaction with NPs and the possible non-B DNA forms detected via CD and transmission electron microscopy (TEM). **b**-**e**, Circular dichroism (CD) spectra showing nanoplastic-induced conformational changes in defined GC DNA duplexes: (**b**) GC tri-repeat duplex (6 bp); (**c**) GC hexa-repeat duplex (12 bp); (**d**) longer GC-rich duplex (25 bp) with extended sequence context; and (**e**) methylated GC hexa-repeat duplex. In all panels, native DNA (dark green) is compared to DNA incubated with polystyrene nanoparticles (PS, blue), aminated polystyrene (PS–NH₂, red), and carboxylated polystyrene (PS– COOH, light green). **f-g**, TEM of the unexposed plasmid control in the presence of anti-DNA Z antibody (**f**) in comparison to form V plasmid upon exposure to NPs (**g**). Plasmid indicated with blue arrows. The topological reorganization is represented by schematics below the images. **h**, Circular CD spectra showing nanoplastic-induced conformational changes in defined pyrimidine-rich DNA duplex TCC tetra-repeat incubated with PS NPs (blue), PS-NH_2_ NPs (red), PS-COOH NPs (light green) and without NPs (green). The charged cytosine indicates the corresponding peak at 280 nm.

We first conducted in-vitro circular dichroism (CD) spectroscopy as a gold standard for assessing DNA secondary-structure transitions^75^. Based on the length and the sequence, B-DNA displays a positive CD peak at ∼ 275nm (π- π base stacking) and a negative peak between 245 and 255nm (sugar–phosphate torsion, χ angle)^75^ (**Fig. S5a**, green spectrum, B-DNA vs blue spectrum, Z-DNA). Then, we tested heteroduplexes of GC triplet repeat (∼2,38nm in length, **Fig. 5b**), hexarepeat (∼4nm in length, **Fig. 5c**), and BZ heteroduplex that contain GC and AT regions (∼8.5nm in length, **Fig. 5d**). Upon interaction with the NPs, T(CG)3 heteroduplex preserved the right-handed helices but the B-form helicity marker shifted the spectra of the DNA-NP complexes from the one typical for the B-DNA (**Fig. 5b**, green) to a spectrum typical for the ѱ+ DNA (**Fig. 5b**, blue, red and light green), which is reported for DNA assemblies into a cholesteric phase^73^. In the extended heteroduplexes, the χ angle shifted from –4° to ≈ 0°, exceeding the intrinsic thermal fluctuation range (2–3°), indicative of distorted B-DNA topology (**Fig. 5c-d**, compare green spectrum versus blue, red and light green). Such changes in the spectra are indicative of preserved right-handed helix, but the collapse of the B-form marker can signify the presence of an unwound intermediate state of atypical A-DNA or Z-DNA. Moreover, this NP-effect appears to be specific to the GC sequences since the AT₆ and AC₆ duplexes exhibited only minor χ angle variations within intrinsic noise (3–4°) (**Fig. S5b-c**, spectrum of untreated DNA in green, treated DNA – in blue).

It has been previously reported that cytosine methylation alters DNA mechanics^76^. Therefore, we also assessed whether methylation of cytosine could impact the interaction between the NPs and DNA. The spectrum of the methylated DNA heteroduplex exhibited more enhanced classical features of B-DNA compared to its unmethylated counterpart (**Fig. S5d**, compare green to blue) confirming that cytosine methylation provides higher stability. Nevertheless, when the methylated DNA was exposed to the NPs, at least in the case of the GC hexarepeat, we did not observe any protective effect of the DNA methylation since the B-DNA marker of the methylated DNA collapsed from -5 to 0 mdeg in the presence of the NPs (**Fig. 5e**, light green to blue and red). Additionally, we tested if the NPs could promote B-to-Z-DNA transitions in conditions, such as 20% ethanol shown to stimulate Z-DNA formation only in methylated DNA but we observed the same effect of the NPs, i.e. reduced negative peak at around 250nm (**Fig. S5e**-**f**). The spectral shifts observed only with GC heteroduplexes (Fig. 5b-e and **Fig. S5e**-**f**) indicate sequence-guided alterations in DNA secondary structure, including features consistent with B–A and B-Z transitions.

Unlike linear DNA, circular DNA sustains torsional stress. Therefore, we next tested whether NPs can trigger global topological inversion of pBR322, shown to contain Z- DNA sequences^77^. While linear DNA can freely dissipate torsional strain through rotation, circular plasmids are topologically constrained and accommodate stress by flipping their helical handedness. In short, even if the entire plasmid does not comprise Z-DNA sequence, only if a substantial fraction of the plasmids adopts left-handed conformation, this can result in the formation of the so-called V-form plasmid, i.e., a highly coiled, duplex circular DNA^77^. CD spectra of pBR322 plasmid incubated with NPs showed a typical B-DNA spectrum while the plasmid incubated in 4M NaCl showed a spectrum typical for C-DNA – left-handed DNA with a B-DNA backbone in which the minor groove is deeper as a result of base-pairs being moved from the center^71,78^ (**Fig. S5g**, compare blue to green). Interestingly, plasmids exposed to the different types of NPs show similar CD spectra to one of the plasmid incubated with 4M NaCl, known to induce B→Z DNA transitions^75^ (**Fig. S5h**), indicating that NPs can promote a right to left (R->L) handedness change in circular DNA that contains Z- promoting sequences. To confirm that the NPs induced R->L transition in the plasmid, we performed TEM of the plasmids in the presence of anti-Z-DNA antibodies incubated with and without PS-NH_2_. Interestingly, while the untreated plasmid did not show any super-coiled structures (**Fig. 5f**, plasmid borders indicated with blue arrows), we detected the V form of pBR322 only in the samples directly exposed to the NPs (**Fig. S5g**, plasmid borders indicated with blue arrows, black spheres on the DNA string indicative for Z-DNA-antibody complex). Moreover, the supercoiled structure was further confirmed after purification of the plasmids from the NPs and antibodies (**Fig. S5i-j**). In conclusion, while native plasmid DNA displays a relaxed, open circular conformation, NP exposure induces pronounced compaction, looping, and irregular contouring, consistent with NP-driven perturbation of DNA topology and consistent with the form V of pBR322 induced by Z sequences shown by Lang et al.^77^. To verify that such interactions can take place *in vivo*, we tested the genomic DNA extracted from strains TA1535 and TA1538 exposed to PS-NH₂. The genomes of the bacteria exposed to the most mutagenic NPs (**Fig. 1b**), also showed attenuated ellipticity and partial unwinding, indicating conformational distortion suggesting R-to-L transitions (**Fig. S5k**).

Since we established a direct sequence- and length-specific non-B-DNA conformational polymorphism, we tested if the observed substitutions in the CCCP-treated bacteria, triggered by PS-COOH NPs, can be because of base protonation, i.e., how heteroduplexes formed by (TC)6 and (TCC)4 repeats are affected by the presence of NPs. The presence of polypyrimidine or polypurine DNA stretches provide the platform for the formation of H-DNA - a triplex-DNA structure in which one strand folds back and binds to an existing heteroduplex via Hoogsteen (non-Watson-Crick) base pairing, the protonation of cytosine (C+) being the hallmark of DNA triplex formed by C-rich polypyrymidine^79^. The spectra of the polypyrimidine heteroduplexes, used in this study, exhibited ѱ+ DNA spectra (**Fig. 5h** and **Fig. S5l-m**, green). The addition of PS-COOH NPs triggered surface-templated Hoogsteen induction, characterized by a positive peak at 288 nm as well as a negative peak at 260 nm (**Fig. S5l**, green vs blue spectrum). To test if the effect of the PS-COOH NPs could be amplified, we repeated the experiment with a TCC tetrarepeat and tested the activity of the PS, PS-NH_2_ and PS-COOH NPs (**Fig. 5h**). While the presence of NPs disrupted the B-conformation negative peak (**Fig. 5h**, blue, red, and light green spectra), the presence of PS-COOH triggered local protonation of the cytosine and displayed the strongest increase of the peak at 280nm (**Fig. 5h**, light green spectrum). Additionally, the presence of PS NPs changed the spectrum of the (TC)6 sequence but it did not show any of the features typical for the formation of H-DNA (**Fig. S5m**, green vs blue spectrum), i.e., a highly positive peak at ∼288nm due to C+ and a negative peak between 250-260nm.

### NPs control the frequency of spontaneous mutations in a DNA methylation manner

The mutation frequency in bacteria is often guided by the underlying DNA methylation^80^. Since we did not observe any structural effect of methylation on NP-DNA interactions, we tested whether DNA methylation may play a role against NP-induced mutagenesis. To do that, we undertook a biochemical approach by performing Ames assays in cells pre-treated with the DNA methyltransferase inhibitor 5-aza-deoxycytidine (5-aza-dC)^54–56^. It should be noted that not only does 5-aza-dC inhibit DNA methylation, but also, being a modified base, it can be integrated into the bacterial chromosome^81^ without affecting bacterial viability (**Fig. S1e**). Additionally, we undertook a genetic approach by testing *Escherichia coli* K-12 wild type and its DNA-methylation deficient mutants for rifampicin resistance – a method used to evaluate the mutation rate of a bacterial strain^82^ (see **Fig. 6a** for experimental design).

**Figure 6:**
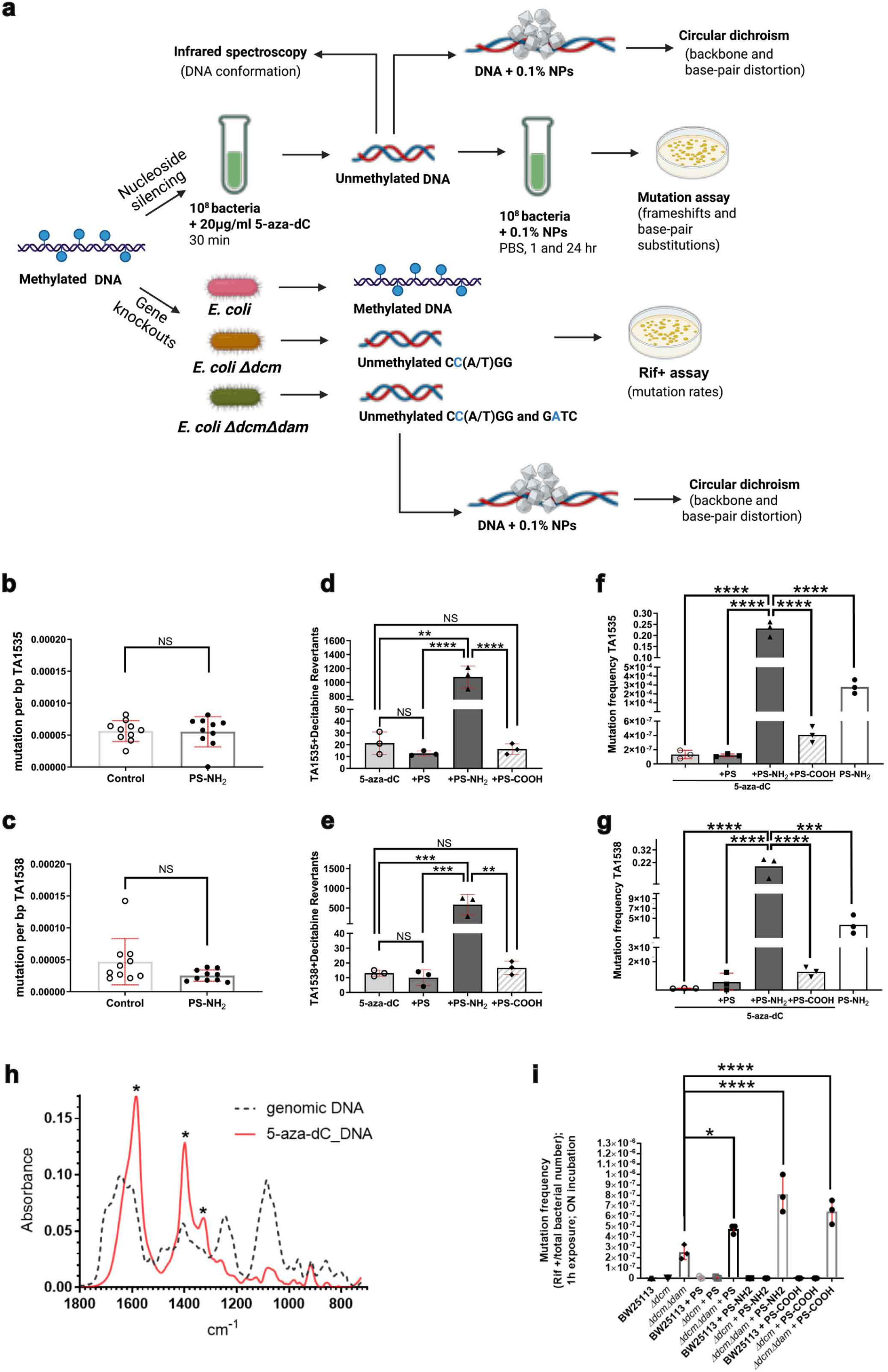
Effects of nanoplastics and epigenetic states on mutational burden and frequency. **a**, Schematic overview of the experimental setup for testing the role of methylation in NP-mediated mutagenesis. The biochemical inhibition of DNA methylation with 5-aza-dC and the genetic approach involving in-frame deletions of the DNA methyltransferase genes are outlined. **b**-**c**, Mutation rate in TA1535 (**b**) and TA1538 (**c**). **d-e**, Ames test quantification of revertants in 5-aza-2′-deoxycytidine (decitabine)-treated bacteria exposed to PS, PS–NH₂, PS–COOH NPs, showing a marked increase in (**d**) base-substitution and (**e**) frameshift revertants after exposure to PS-NH₂. **f-g**, Mutation frequencies in decitabine-treated TA1535 (**f**) and TA1538 (**g**) following exposure to nanoplastics with distinct surface chemistries (PS, PS–NH₂, PS– COOH). **h**, Infrared spectroscopic analysis of native genomic DNA and genomic DNA from bacteria incubated with 5-aza-2′-deoxycytidine (decitabine), x-wavelength (cm^-1^) and y-absorbance. **i,** Mutation frequencies in *E. coli* epigenetic mutant strains after 1 h exposure to PS, PS-COOH or PS-NH₂, demonstrating that genetic disruption of DNA methylation pathways sensitizes bacteria to nanoplastic-induced mutagenesis.

Based on the WGS experiments, the PS-NH_2_ nanoparticles did not affect the mutation rate of the tested strains (**Fig. 6b-c**). First, to directly assess how methylation loss influences DNA structure, we recorded CD spectra of chromosomes from 5-aza-dC-treated bacteria. CD spectra of chromosomes from 5-aza-dC-treated bacteria displayed a B-DNA spectrum (**Fig. S6a**, green curve), and addition of PS-NH_2_, shifts the 260 nm peak to 275 nm, which is indicative for altered base-stacking interactions (**Fig. S6a**, blue curve). Then we tested if the incubation of bacteria with 5-aza-dC alone had any effect on spontaneous mutation frequency, and as displayed in **Fig. 6f-g**, we found that the effects are negligible. It is important to highlight that the cell number of the 5-aza-dC-treated bacteria prior NP exposure remained the same (**Fig. S1e** samples TA1535+5-aza-dC and TA1538+5-aza-dC). Yet, when exposed to NPs, the presence of PS-NH₂ NPs simultaneously reduced bacterial viability 100 times (**Fig. S6b**) and generated a similar number of revertants to the revertants obtained after exposing the untreated bacteria to PS-NH₂ NPs (**Fig. 6d-e**), revealing that methylation, at least partially, provides functional protection against NP-induced mutagenesis. This drastically increased the mutation frequencies of TA1535 and TA1538, i.e. 10^5^ in comparison to the spontaneous revertants and 10^3^ more compared to the PS-NH_2_- treated bacteria (**Fig. 6d-e**). Similar but less powerful effect was exerted by the PS-COOH NPs on CCCP treated TA1535 bacteria as the mutation frequency of the CCCP-treated bacteria exposed to PS-COOH increased 6 times in comparison to the spontaneous revertants of the CCCP-treated bacteria and it was 20 times higher than the revertants generated by the TA1535 bacteria exposed to PS-COOH (**Fig. S6c**). Taken together, there is a combined effect of methylation loss and NP-induced torsional strain that NPs exert on mutation frequency.

Then, to analyse if the incorporation of 5-aza-dC in the bacterial chromosome has a structural effect on the DNA double helix, we performed IR spectroscopy (**Fig. 6h**). The IR spectrum of the genome extracted from bacteria treated with 5-aza-dC displays distinct features indicating facilitation of R-to-L transition of the bacterial genome upon incubation of the bacteria with 5-aza-dC. The most significant peaks are indicated with stars in **Fig. 6h**, red spectrum. The peak at ∼ 1590 cm^-1^ is assigned to the vibrations of single-stranded dG residues due to base unstacking; the one between 1390-1410 cm^-1^ - to the C3’-endo deoxyribose of deoxyadenine in A- and Z-DNA) and the one at ∼ 1320 cm^-1^ corresponds to the presence of dG in syn conformation in Z-DNA^83^. The fact that there is a decrease in the peaks corresponding to asymmetric and symmetric PO_2_^-^ groups (1240 and 1100 cm^-1^, respectively) further suggests that the incorporation of 5-aza-dC and lack of methylation cause overall conformational changes typical for non-B-DNA forms. To confirm the role of methylation genetically, we examined *Escherichia coli* K-12 wild type and its DNA-methylation deficient mutants: *Δdcm* and *ΔdamΔdcm*. While all genomes displayed similar native CD spectra (**Fig. S6d**, box I), PS-NH₂ exposure consistently reduced the positive CD peak across genotypes (box II -IV), indicating decreased base stacking and duplex rigidity. Correspondingly, the *ΔdamΔdcm* double mutant exhibited up to 2-fold higher mutation frequency 1h of NP exposure -notably with PS-COOH and PS-NH_2_ but also with PS - without affecting viability (**Fig. 6i** and **Fig. S6e**).

## Conclusion

Attempts to investigate NP-induced genotoxicity have been made globally and with different model systems – from cells, tissues and organs to a variety of biological species^51,84,85^. The potential mechanisms for NP-induced genotoxicity are generally attributed to oxidative stress and indirect effects on DNA repair pathways, all of which resulted either in cell cycle arrest, apoptosis or pathological outcomes, such as endothelial leakage, dysbiosis and epithelial-to-mesenchymal transition in lung tissues^51,84,85^. Even though little evidence suggests that some NPs, especially PS, may be genotoxic, the presence of NP-induced mutagenesis has remained a much bigger and unsettled question.

Here we report a direct and underappreciated mechanism of mutagenesis, i.e., physical interaction of nanoparticles with DNA that alters its topological state, conformational stability, and hence mutation frequency. This interaction is guided by GC-sequence specific polymorphism that can promote formation of non-B-DNA structures such as atypical Z-DNA or H-DNA. Across five experimental tiers—ranging from bacterial mutagenesis and whole-genome spectra (**Fig. 1b**, **Fig. 2**, **Fig. 4g** and **Fig. 6**), nanoparticle uptake (**Fig. 1c-d**, **Fig. 3** and **Fig. 4**), to direct topological analysis (**Fig. 3d-e** and **Fig. 5**) — polystyrene nanoplastics exert their mutagenic potential based on their surface chemistry and the physiological state of the bacteria.

When tested for mutagenicity, NPs can be viewed as conditional mutagens. For example, PS-NH_2_ is either mutagenic (by triggering frameshifts and base substitutions) or toxic. Thus, it acts by physically generating non-B-DNA species without necessarily unwinding the DNA heteroduplex. This mutagenicity is further enhanced by the fact that previous work on mammalian and plant systems has shown that PS nanoparticles—particularly cationic variants—can cause, as previously reported, DNA double-strand breaks, chromosome rearrangement, and glutathione depletion without requiring nuclear entry ^51,52,86^. On the other hand, PS-COOH NPs are a conditional mutagen because they are only mutagenic under specific environmental conditions that provide proton uncoupling which results only in base substitutions (**Fig. 4g** and **Fig. S4d**). Based on our in-vitro studies (**Fig. 5h**), the presence of PS-COOH NPs causes local protonation of the cytosine (C+) in a polypurine.polypyrimidine sequence-specific manner, without disrupting the global helical structure. So far, C+ has been associated predominantly with point mutations^87,88^, consistent with the base substitution effect observed when bacteria were incubated with CCCP and exposed to the NPs.

To extend this concept to the two primary forms of genetic material, i.e. circular and linear DNA, we demonstrated *in vitro* that PS-NPs directly disrupt B-DNA architecture and promote non-B conformers, including Z-like DNA in plasmids, which are known to favour frameshift and base substitution mutations, and alternative A- and H-like forms. Such non-canonical structures are known mutational hotspots that promote frameshifts and base substitutions via polymerase slippage or misalignment. The specific GC sequence dependence we observed, is consistent with prior in-vitro studies showing that cationic NP (albeit non-plastic: gold and poly(amidoamine) dendrimers) charge and geometry can modulate duplex stability, minor groove hydration, and local bending stiffness^54^. In-vitro spectroscopy and TEM confirmed that PS–NH₂ nanoparticles induce a helical flip of supercoiled plasmid DNA (**Fig. 5g**), converting right-handed B-DNA to left-handed Z-like DNA. This topological stress, detectable as a mirror-flipped CD signature, parallels the effects of cationic dendrimers or polylysine reported compacting or unwind duplexes ^39,40^.

Additionally, loss of methylation via 5-azadeoxycytidine or genetic deletion of *dam/dcm* methyltransferases sensitized bacterial DNA to PS–NH₂, enhancing mutagenicity and weakening base-stacking interactions (**Fig. 6**). This protection likely stems from methylation-induced increases in DNA stiffness and helical persistence length, properties known to resist non-B structural transitions, which reduces susceptibility to non-canonical structural transitions in both prokaryotic and eukaryotic genomes^54^. Together, these results connect NP surface chemistry to mutational mechanisms through DNA topology. Aminated PS–NPs, by virtue of their positive surface charge, electrostatically couple to the DNA backbone, altering superhelicity, bending stiffness, and local hydration—each a determinant of helical handedness^48^. The resulting B→Z flips or partial unwinding events generate transient non-B intermediates that serve as templates for frameshifts and base substitutions, consistent with the distinct GC motifs revealed by whole-genome sequencing and in-vitro CD experiments. This topology-driven mutagenic mechanism redefines how NP may impact genomic stability. While oxidative stress and DNA repair interference remain relevant, our data indicate that electrostatic and conformational effects alone are sufficient to generate mutations—a mode of action previously observed in systems where cationic nanoparticles bind and compact DNA via electrostatic interactions, altering its higher-order structure^89^.

In short, this study shows that nanoplastics are not mutagens in a simple yes-or-no sense. In bacteria, selected nanoplastics can enter living cells, associate with the chromosome and disturb the normal structure of DNA, creating conditions in which mutations can arise. Whether this happens—and whether exposure results in mutation, delayed mutation, cell death or no detectable effect—depends both on the properties of the particle and on the state of the cell and its genome. We therefore propose that nanoplastics can act as conditional mutagens: their mutagenic effects emerge only when a particular nanoparticle encounters a susceptible cellular and genomic state. We propose the term “conditional mutagen” to describe an agent whose mutagenic outcome is not determined by the agent alone, but emerges from the interaction between its physicochemical properties and the physiological and genomic state of the exposed cell. By this definition, nanoplastics can act as conditional mutagens: particle size, surface and backbone chemistry determine genome access, whereas cellular energetics and DNA methylation influence whether that interaction results in mutation, delayed mutation or cell death.

## Acknowledgements

We acknowledge the Department of Molecular Biology, Umeå University Sweden for providing laboratory space and financial support and the Head of the department, Prof. Matthew Francis, for organizational support specifically. We acknowledge the facilities and technical assistance of Vibrational Spectroscopy Core Facility (ViSp) and the BioMolecular Characterization Umeå (BMCU), Umeå University, Sweden. We also acknowledge the CF Genomics and the CF Bioinformatics supported by the NCMG research infrastructure (LM2023067 funded by MEYS CR) for their support in obtaining the scientific data presented in this paper. We thank Anton Jäger from the Department of Pathology, Medical University of Vienna, for support in graphical artwork. L.K. acknowledges the support from MicroONE, a COMET Modul under the lead of CBmed GmbH, which is funded by the federal ministries BMK and BMDW, the provinces of Styria and Vienna, and managed by the Austrian Research Promotion Agency (FFG) within the COMET—Competence Centers for Excellent Technologies—program. Financial support was also received from the Austrian Science Fund (grants FWF: P26011, P29251, P 34781), the Vienna Science and Technology Fund (WWTF), grant number LS19-018.

## Credits

1) Conceptualization/Planning, 2) Methodology, 3) Formal analysis, 4) Experimental work, 5) Original draft, 6) Reviewing and finalizing draft, 7) Supervision, 8) Data curation, 9) Visualisation, 10) Project administration, 11) Funding acquisition: N.Z.:1,2,3,4,5,6,7,8,9,10; I.K.: 2,3,4,8,9; F.M.: 1,2,3,8,9; L.M.: 2,3,4,5,9; J.O.: 1,2,3,4; R.L.: 4,9; A.M.: 2,3,4; M.P.: 7; V.B.: 2,3,8; UR: 1,2,3,5,6,9; L.K.: 1,2, 5,6,7,10,11.

## Abbreviaions

NPs: nanoplastics
PS: Polystyrol
PS-NH2: aminated polystyrol
PS-COOH: carboxylated polystyrol
PMMA: poly-(methyl methacrylate)
LB: lysogeny broth

## Supplementary Figures

**Figure S1:**
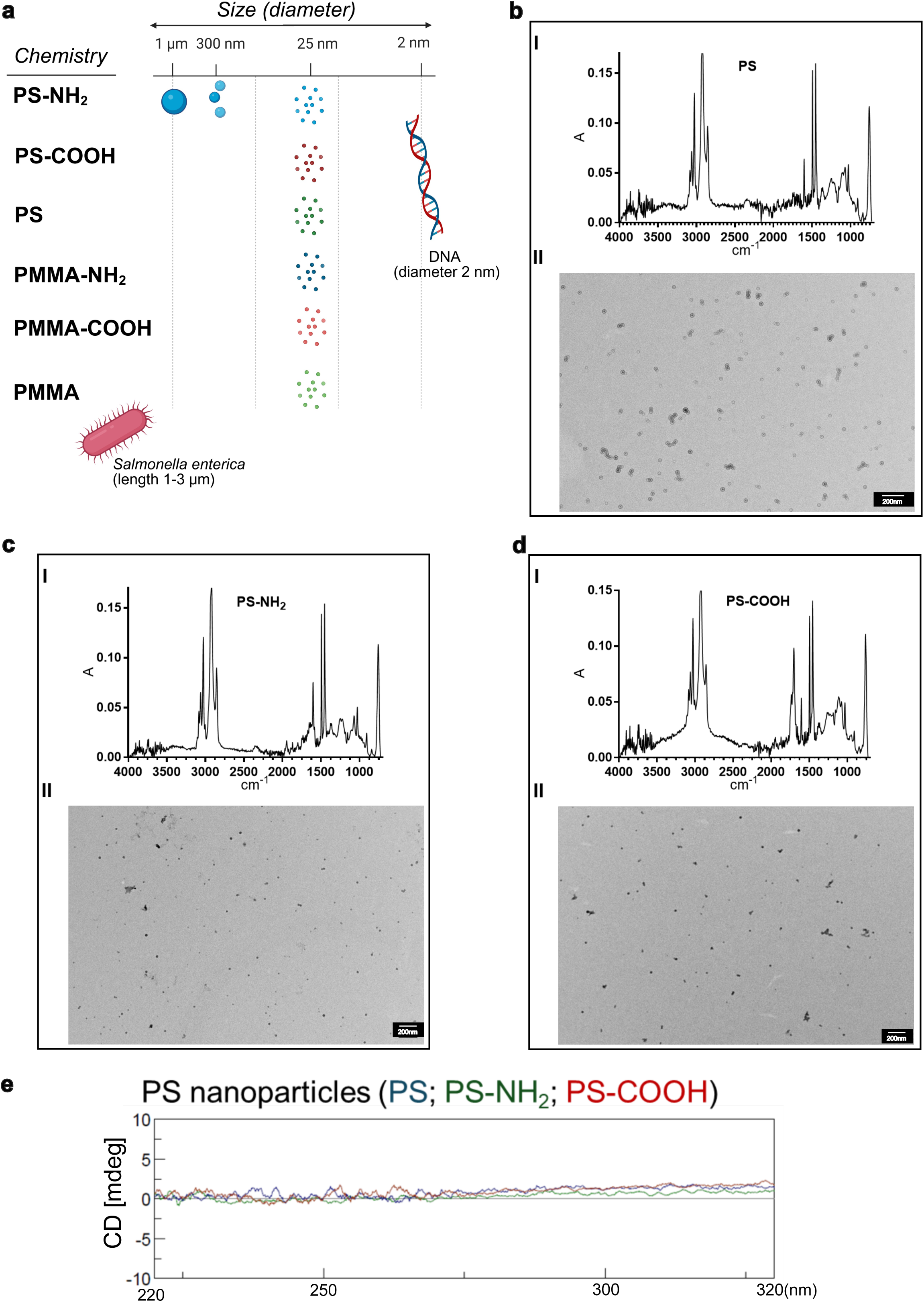
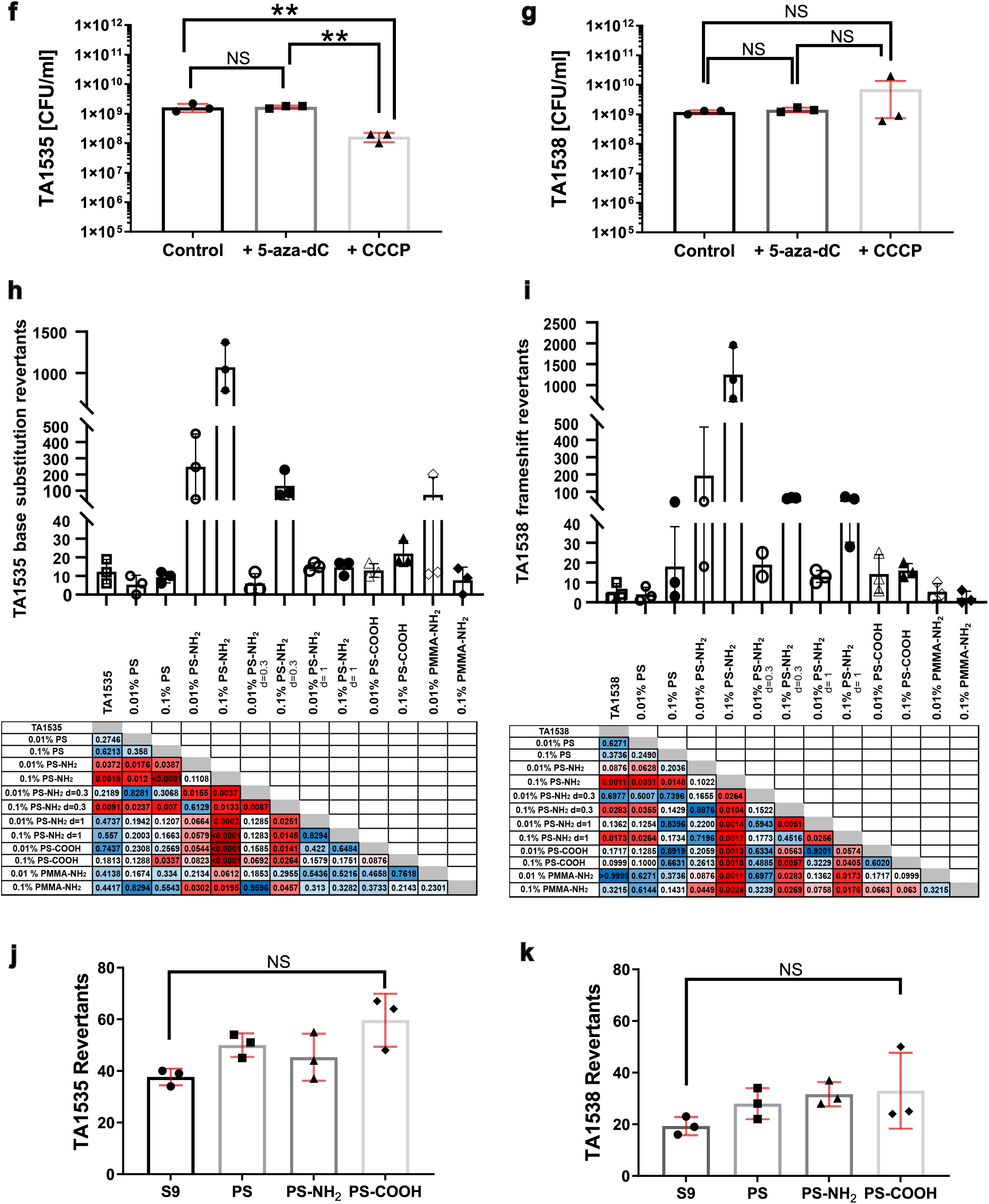
Physicochemical characterisation of polystyrene (PS) nanoplastics and control experiments used in mutagenicity assays. **a**, Surface chemistry and physical properties of PS particles used in this study in connection to bacterial cells and DNA. The nanoplastic PS is represented by nanospheres with d = 25nm with three surface chemistries - non-modified (PS), carboxylated (PS-COOH) and aminated (PS-NH_2_) nanoplastics. To distinguish how NP physical size and surface chemistries affect biological systems, aminated (PS-NH_2_) nanospheres with d = 0.3µm and 1µm, as well as modified and non-modified polymethyl methacrylate (PMMA) nanoparticles with with d = 25nm are employed as controls. **b**–**d**, Infrared spectroscopic imaging (I) and transmission electron microscopy (TEM) (II) of PS (**b**), PS–NH₂ (**c**), and PS–COOH (**d**) nanoplastics. Spectra display characteristic absorbance peaks (x axis = wavelength cm⁻¹; y axis = absorbance), and TEM images show particle morphology and size distribution. Scale bar, 200 nm. **e**, Circular dichroism (CD) spectra of PS, PS–NH₂, and PS–COOH NPs showing minimal signal across the measured range, consistent with the non-chiral nature of PS-NPs. **f-g**, Bacterial cell counts of strain TA1535 (**f**) and TA1538 (**g**) bacteria before NP exposure, confirming comparable inoculum sizes across all treatment groups used in uptake and mutagenicity experiments. **h-i**, Nanoplastic-mediated mutagenesis of strain TA1535 (**h**) and TA1538 (**i**) after exposure to 0.01% and 0.1% (w/v) PS, PS–NH₂, PS-COOH, PMMA, PMMA–NH₂, and PMMA-COOH nanoparticles of 25-nm diameter for 1 hour; PS–NH₂ with 0.3µm and 1µm diameter micro-nanoparticles were included as controls. Y-axis indicates the strain and the type of mutation it detects. **j-k**, Co-incubation with 2% S9 liver extract abolished NP-induced mutagenesis, returning revertant counts to baseline levels of TA1535 (**e**) and TA1538 (**f**) bacteria.

**Figure S2:**
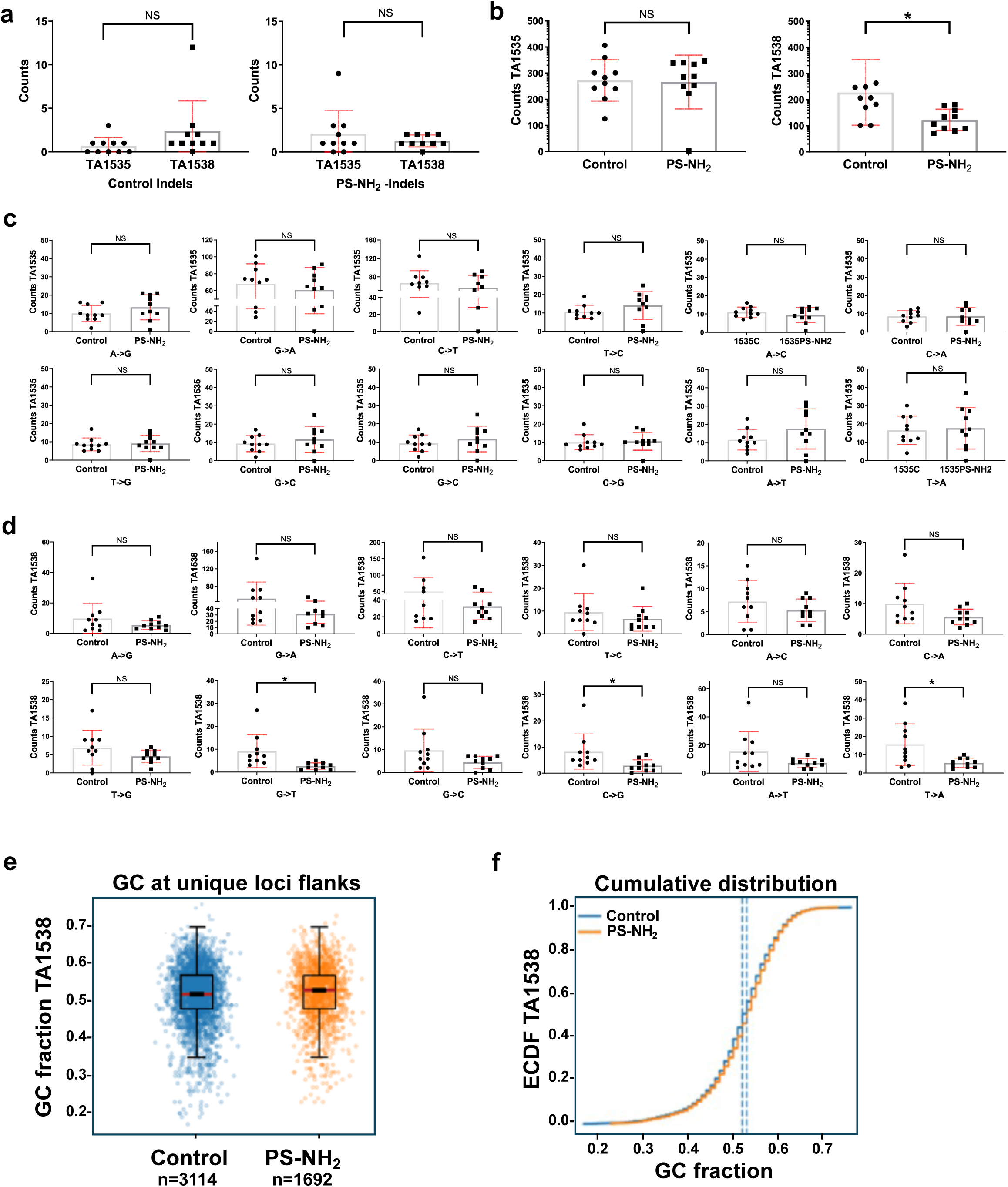
Mutational analysis of the spontaneous and NP-induced revertants. a, Indel spectra of TA1535C and TA1535PS–NH₂ as well as of TA1538C and TA1538PS–NH₂. **b**, Unique base-substitution spectra of TA1535C and TA1535PS– NH₂ as well as of TA1538C and TA1538PS–NH₂. **c**, Base-substitution spectra of TA1535C and TA1535PS–NH₂. **d**, Base-substitution spectra of TA1538C and TA1538PS–NH₂. **e**, CG fraction at the unique sites of mutations in spontaneous TA1538C (blue) and NP induced TA1538PS–NH₂ (orange) with 100 nucleotids upstream (-) and 100 nucleotides downstream (+) combined (MWU -Mann-Whitney U). **f**, Empirical cumulative distribution functions (ECDFs) of the GC content in ±100 bp flanking regions in spontaneous TA1538C (blue) and NP-induced TA1538PS–NH₂ (orange) revertants.

**Figure S3:**
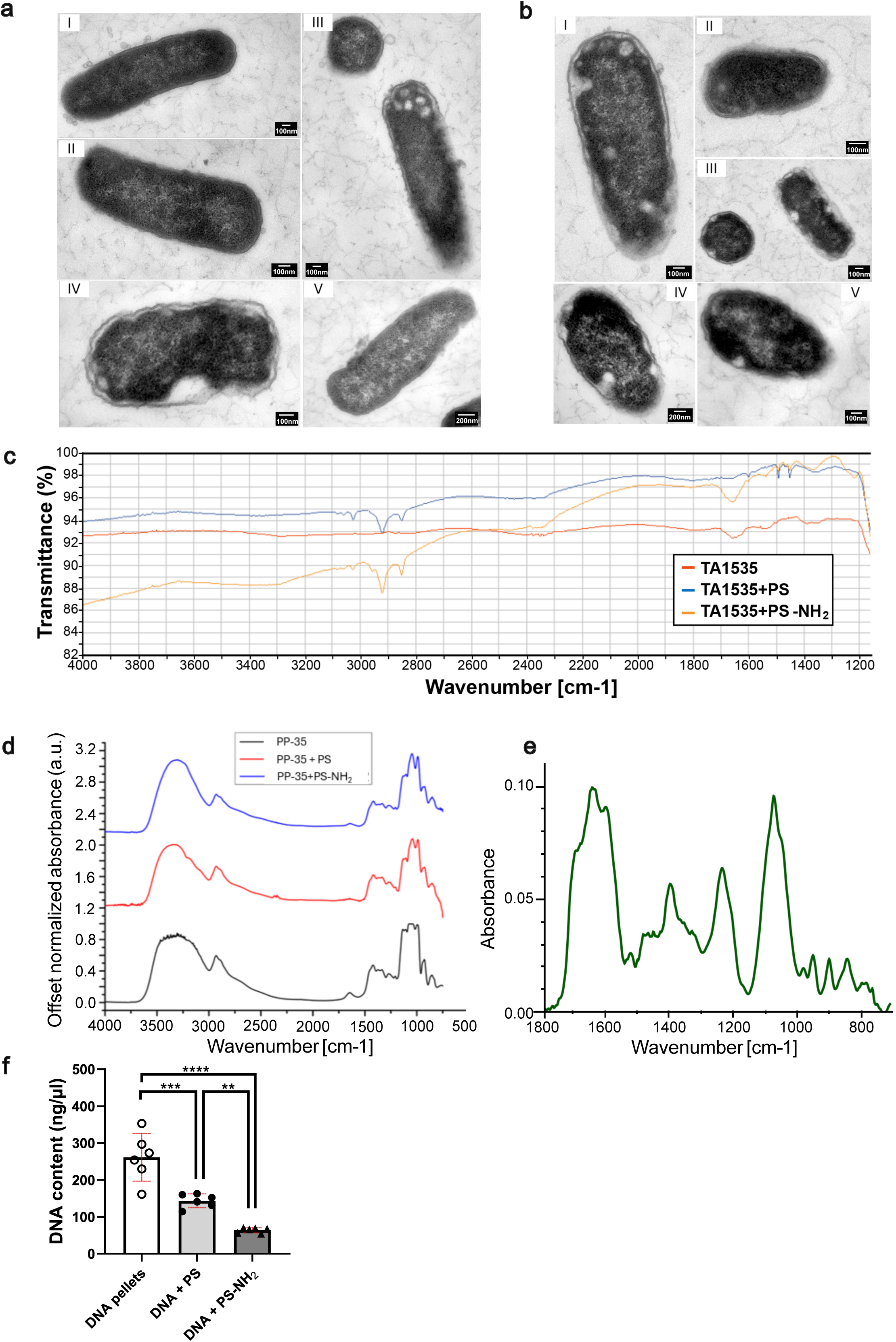
Infrared spectroscopy bacterial DNA-NP complexes. **a**-**b**, Ultrastructural imaging of *Salmonella* TA1535 (boxes I, II and III) and TA538 (IV and V) with PS (**a**) and PS-COOH (**b**). **c**-**d**, IR spectroscopy of different bacterial compartments. **c,** IR spectra of the chemical environment of strain TA1535 (designated as S for supernatant) in the absence of NPs (red) and in the presence of PS (in blue) and PS-NH_2_ (in orange). **d,** Periplasmic fractions of strain TA1535 unexposed to NPs (in black), exposed to PS (in red) and to PS-NH_2_ (in blue). **e,** Canonical DNA vibrational signatures displayed by the bacterial chromosome as a reference for intracellular biomolecular composition. **f**, Quantification of DNA release from the DNA-enriched pellets from bacteria exposed to the plastics.

**Figure S4:**
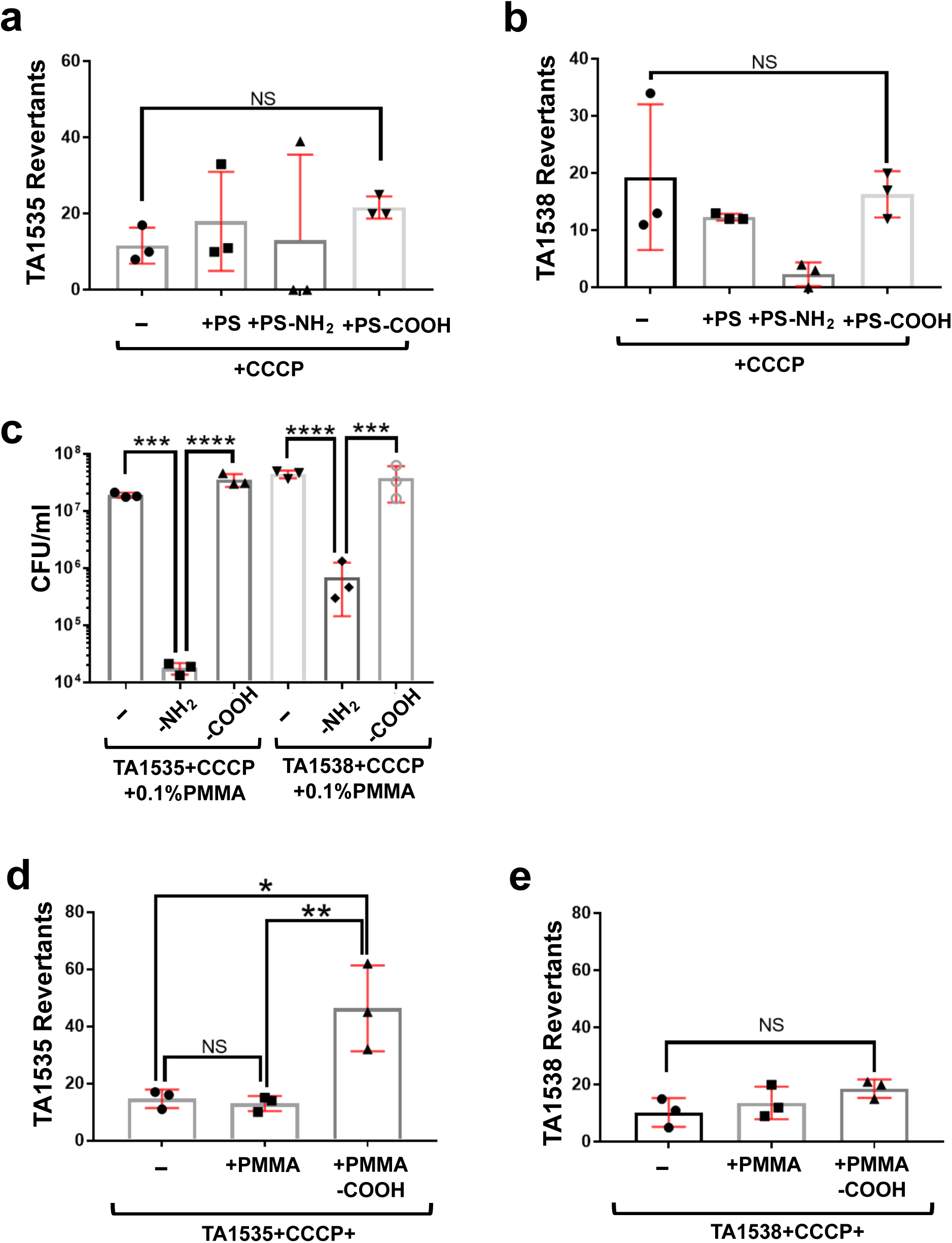
The role of membrane potential and surface chemistry of NPs in mutagenic outcomes in *Salmonella enterica* TA1535 and TA1538. a-b, Ames reversion assay of CCCP-treated cells exposed to PS-NPs for 1h with TA1535 (a) and TA1538 (b). c, Viable cell counts (CFU ml⁻¹) of CCCP-treated bacteria exposed to 0.1% PMMA (TA1535+CCCP 0.1% PMMA and TA1538+CCCP 0.1% PMMA), 0.1% PMMA–NH₂ (TA1535+CCCP 0.1% PMMA–NH₂ and TA1538+CCCP 0.1% PMMA– NH₂) and 0.1% PMMA-COOH (TA1535+CCCP 0.1% PMMA-COOH and TA1538+CCCP 0.1% PMMA-COOH) after exposure for 24h. **d**, Ames reversion assay of CCCP-treated TA1535 cells exposed to 0.1% PMMA, PMMA–NH₂ and PMMA-COOH NPs overnight. **e**, Ames reversion assay of CCCP-treated TA1538 cells exposed to 0.1% PMMA, PMMA–NH₂ and PMMA-COOH NPs overnight.

**Figure S5:**
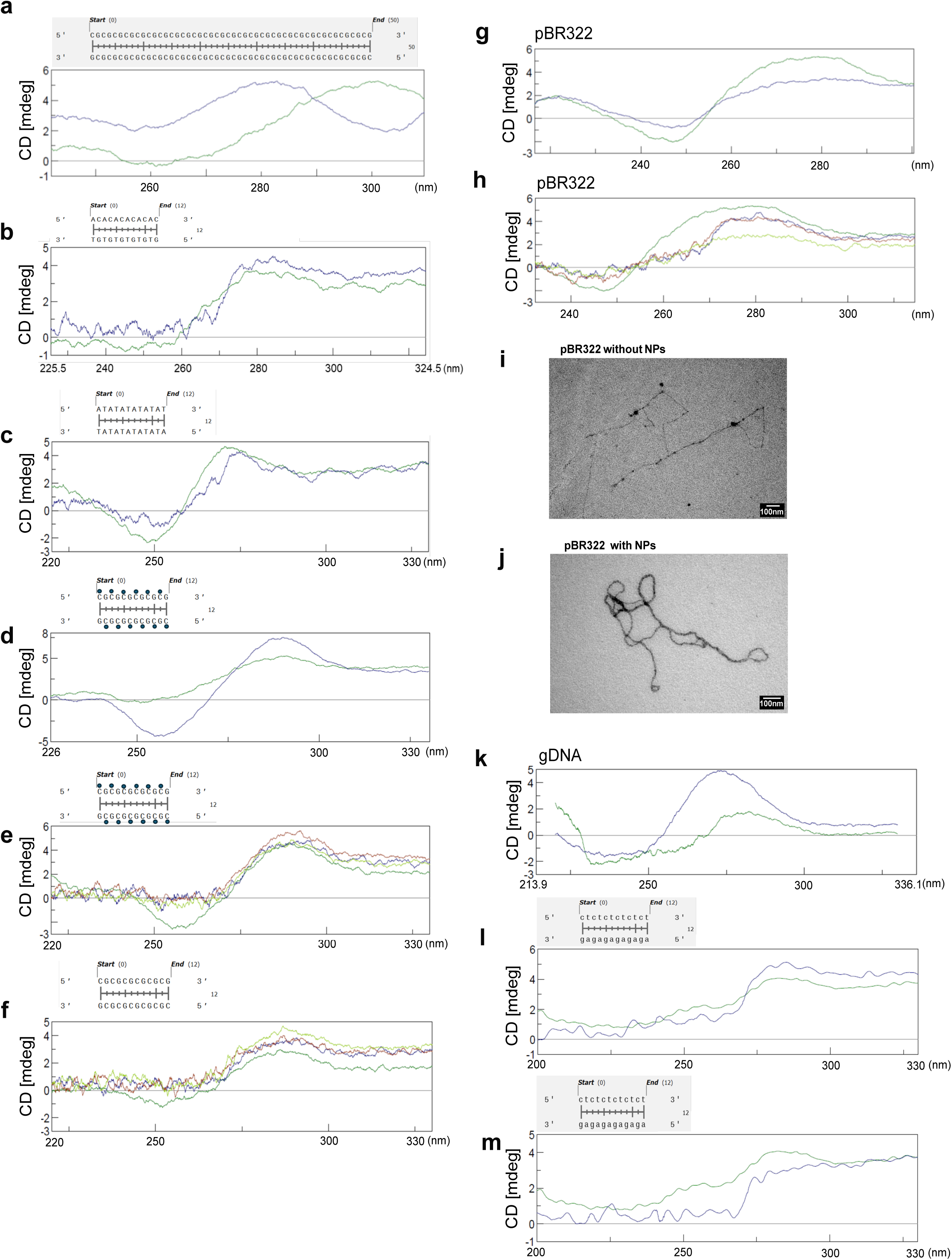
Circular dichroism (CD) analysis of NP–DNA interactions and methylation effects. **a**, CD spectra of B- (green) and Z-DNA (in blue) of 25-nt CG repeat. **b**,**c** CD spectra of AC (**b**) and AT repeat (**c**) duplexes (12 bp) interacting with PS-NH₂ in the absence (green) and presence of aminated polystyrene nanoparticles (PS–NH₂, blue), showing sequence-dependent susceptibility to NP-induced structural perturbation. **d**, CD spectra of methylated (blue) and unmethylated (green) CG hexarepeat (12 bp). **e**-**f**, CD spectra of methylated (**e**) and unmethylated (**f**) CG hexarepeat (12 bp) in 20% ethanol. Native DNA (green) is compared with DNA incubated with polystyrene nanoparticles (PS, blue), aminated polystyrene (PS–NH₂, red), and carboxylated polystyrene (PS– COOH, light green), revealing methylation-dependent modulation of nanoplastic-induced conformational changes. **g**-**j**, Influence of Z-prone sequences on plasmid topology. **g**, CD spectra of pBR322 plasmid DNA (green) and plasmid in 4M NaCl (blue). **h**, CD spectra of pBR322 plasmid DNA (green), plasmid + PS (blue), plasmid + PS-NH₂ (orange) and plasmid + PS-COOH (light green), demonstrating PS-NP-induced topological and conformational changes resembling Z-DNA-like signatures; TEM of the unexposed plasmid control in the presence of anti-DNA Z antibody (**g**) in comparison to the V form plasmid upon exposure to NPs (**h**) after antibody and NP purification. **k**, CD spectra of genomic DNA (gDNA) from *Salmonella enterica* in the absence (blue) and presence (green) of 0.1% PS–NH₂. **l**, CT hexarepeat incubated with PS NPs (blue) and without PS NPs (green). **m**, CT hexarepeat incubated with PS-COOH NPs (blue) and without PS-COOH NPs (green).

**Figure S6:**
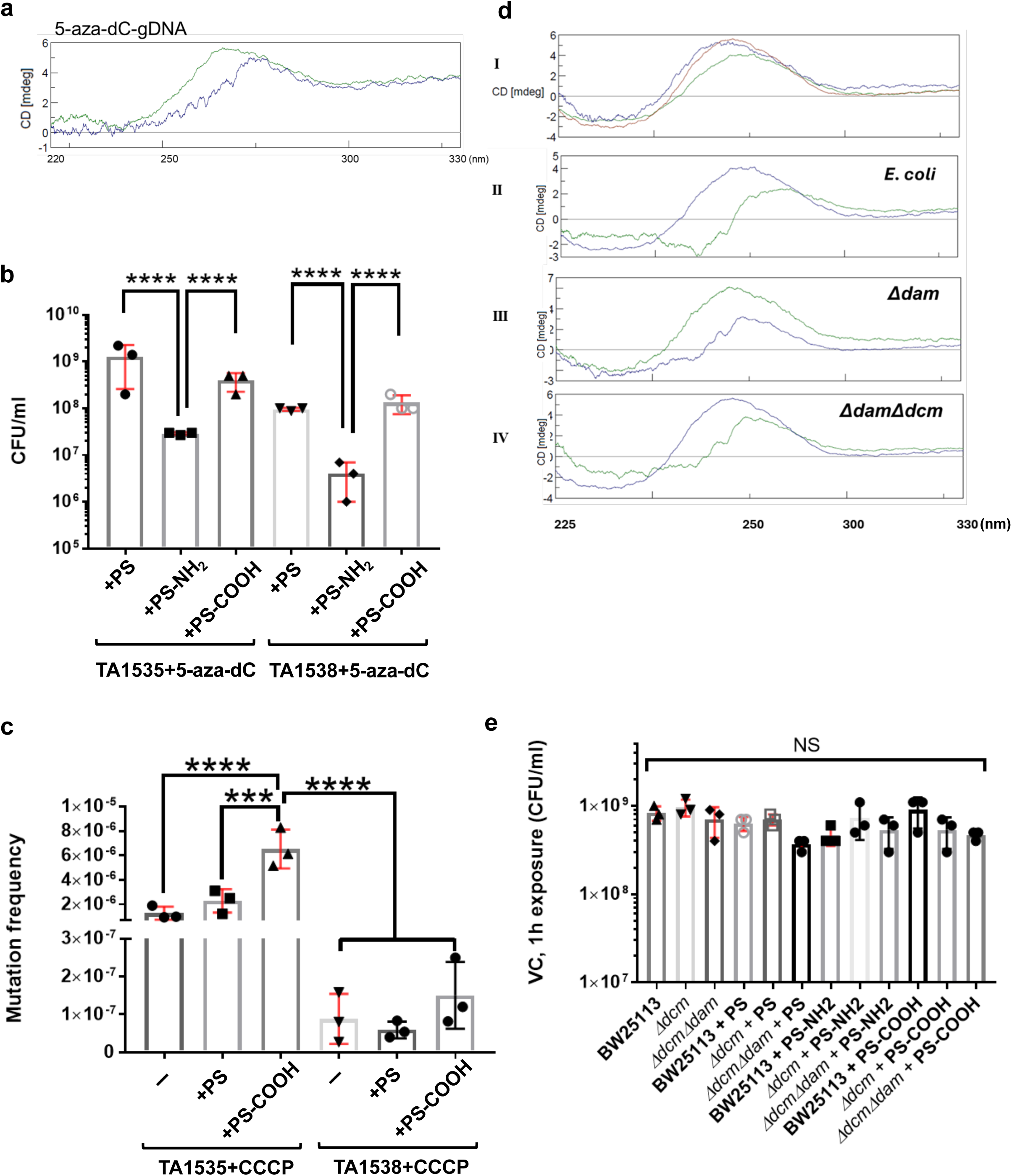
Nanoplastic-DNA interactions related to epigenetic state. **a**, CD spectra of chromosomes from decitabine treated bacteria before (green) and after exposure (blue) to NPs, showing a shift from B- to a non-canonical A-like conformation under hypomethylated conditions **b**, Viable cell counts of decitabine treated bacteria after exposure to NPs with different surface chemistries, indicating 100 times reduction of bacterial viability after PS-NH₂ exposure. **c**, Mutation frequencies in CCCP-treated TA1535 following exposure to PS–COOH, which induced mutation frequencies ∼6-fold higher than CCCP-treated controls and ∼20-fold higher than PS–COOH exposure alone. **d**, CD spectra of genomic DNA from *E. coli* K-12 (green), *Δdam* (blue), and *ΔdamΔdcm* (red). ii–iv, CD spectra of untreated DNA (blue in II and III, and green in III) and 0.1 % PS-NH₂-exposed DNA (complementary color) from K-12 (ii), *Δdam* (iii), and *ΔdamΔdcm* (iv), showing consistent reduction in positive CD amplitude and spectral shape across epigenetic backgrounds. **e**, Viable cell counts of *E. coli* strains following 1 h nanoplastic exposure to nanoplastics.

## References

1. Rasmussen, S. C. From Parkesine to Celluloid: The Birth of Organic Plastics. Angewandte Chemie International Edition 60, 8012–8016, 10.1002/anie.202015095 (2021).

2. Stubbins, A., Law, K. L., Muñoz, S. E., Bianchi, T. S. & Zhu, L. Plastics in the Earth system. Science 373, 51–55, doi:doi:10.1126/science.abb0354 (2021).

3. Dris, R., Gasperi, J., Saad, M., Mirande, C. & Tassin, B. Synthetic fibers in atmospheric fallout: A source of microplastics in the environment? Marine Pollution Bulletin 104, 290–293, 10.1016/j.marpolbul.2016.01.006 (2016).

4. Oliveri Conti, G., et al. Micro- and nano-plastics in edible fruit and vegetables. The first diet risks assessment for the general population. Environmental Research 187, 109677, 10.1016/j.envres.2020.109677 (2020).

5. Peng, L. et al. Micro- and nano-plastics in marine environment: Source, distribution and threats — A review. Science of The Total Environment 698, 134254, 10.1016/j.scitotenv.2019.134254 (2020).

6. Marfella, R., et al. Microplastics and Nanoplastics in Atheromas and Cardiovascular Events. New England Journal of Medicine 390, 900–910, doi:10.1056/NEJMoa2309822 (2024).

7. Li, T. et al. Micro- and nanoplastics in soil: Linking sources to damage on soil ecosystem services in life cycle assessment. Science of The Total Environment 904, 166925, 10.1016/j.scitotenv.2023.166925 (2023).

8. Gigault, J. et al. Nanoplastics are neither microplastics nor engineered nanoparticles. Nature Nanotechnology 16, 501–507, doi:10.1038/s41565-021-00886-4 (2021).

9. Sun, H., Jiao, R. & Wang, D. The difference of aggregation mechanism between microplastics and nanoplastics: Role of Brownian motion and structural layer force. Environmental Pollution 268, 115942, 10.1016/j.envpol.2020.115942 (2021).

10. Shupe, H. J., Boenisch, K. M., Harper, B. J., Brander, S. M. & Harper, S. L. Effect of Nanoplastic Type and Surface Chemistry on Particle Agglomeration over a Salinity Gradient. Environ Toxicol Chem 40, 1822–1828, doi:10.1002/etc.5030 (2021).

11. Hu, T. et al. Abundance of microplastics and nanoplastics in urban atmosphere. Sci Adv 12, eadz7779, doi:10.1126/sciadv.adz7779 (2026).

12. Moon, S. et al. Direct observation and identification of nanoplastics in ocean water. Sci Adv 10, eadh1675, doi:10.1126/sciadv.adh1675 (2024).

13. Kwon, B. G. et al. Regional distribution of styrene analogues generated from polystyrene degradation along the coastlines of the North-East Pacific Ocean and Hawaii. Environ Pollut 188, 45–49, doi:10.1016/j.envpol.2014.01.019 (2014).

14. Ten Hietbrink, S., Materić, D., Holzinger, R., Groeskamp, S. & Niemann, H. Nanoplastic concentrations across the North Atlantic. Nature 643, 412–416, doi:10.1038/s41586-025-09218-1 (2025).

15. Dimante-Deimantovica, I. et al. Downward migrating microplastics in lake sediments are a tricky indicator for the onset of the Anthropocene. Sci Adv 10, eadi8136, doi:10.1126/sciadv.adi8136 (2024).

16. ter Halle, A. & Ghiglione, J. F. Nanoplastics: A Complex, Polluting Terra Incognita. Environmental Science & Technology 55, 14466–14469, doi:10.1021/acs.est.1c04142 (2021).

17. Commission, E. & Environment, D.-G. f. Nanoplastics – State of knowledge and environmental and human health impacts. (Publications Office of the European Union, 2023).

18. Vert, M. et al. Terminology for biorelated polymers and applications (IUPAC Recommendations 2012). Pure and Applied Chemistry 84, 377–410, doi:doi:10.1351/PAC-REC-10-12-04 (2012).

19. Zhu, L., Zhao, S., Bittar, T. B., Stubbins, A. & Li, D. Photochemical dissolution of buoyant microplastics to dissolved organic carbon: Rates and microbial impacts. J Hazard Mater 383, 121065, doi:10.1016/j.jhazmat.2019.121065 (2020).

20. Corbett, L. N. & Amaral-Zettler, L. A. in Plastics and the Ocean 389–428 (2022).

21. Dawson, A. L. et al. Turning microplastics into nanoplastics through digestive fragmentation by Antarctic krill. Nature Communications 9, 1001, doi:10.1038/s41467-018-03465-9 (2018).

22. Banaei, G. et al. Teabag-derived micro/nanoplastics (true-to-life MNPLs) as a surrogate for real-life exposure scenarios. Chemosphere 368, 143736, doi:10.1016/j.chemosphere.2024.143736 (2024).

23. Qian, N. et al. Rapid single-particle chemical imaging of nanoplastics by SRS microscopy. Proc Natl Acad Sci U S A 121, e2300582121, doi:10.1073/pnas.2300582121 (2024).

24. Lambert, S. & Wagner, M. Characterisation of nanoplastics during the degradation of polystyrene. Chemosphere 145, 265–268, doi:10.1016/j.chemosphere.2015.11.078 (2016).

25. Mendez, N. F. et al. Mechanism of quiescent nanoplastic formation from semicrystalline polymers. Nat Commun 16, 3051, doi:10.1038/s41467-025-58233-3 (2025).

26. Hussain, K. A. et al. Assessing the Release of Microplastics and Nanoplastics from Plastic Containers and Reusable Food Pouches: Implications for Human Health. Environ Sci Technol 57, 9782–9792, doi:10.1021/acs.est.3c01942 (2023).

27. Born, M. P. & Brüll, C. From model to nature - A review on the transferability of marine (micro-) plastic fragmentation studies. Sci Total Environ 811, 151389, doi:10.1016/j.scitotenv.2021.151389 (2022).

28. Blasco, E., Sims, M. B., Goldmann, A. S., Sumerlin, B. S. & Barner-Kowollik, C. 50th Anniversary Perspective: Polymer Functionalization. Macromolecules 50, 5215–5252, doi:10.1021/acs.macromol.7b00465 (2017).

29. Moulay, S. Functionalized Polystyrene and Polystyrene-Containing Material Platforms for Various Applications. Polymer-Plastics Technology and Engineering 57, 1045–1092, doi:10.1080/03602559.2017.1370109 (2018).

30. Durmaz, E. N. et al. Polyelectrolytes as Building Blocks for Next-Generation Membranes with Advanced Functionalities. ACS Applied Polymer Materials 3, 4347–4374, doi:10.1021/acsapm.1c00654 (2021).

31. Li, C., Yan, G., Dong, Z., Zhang, G. & Zhang, F. Upcycling waste commodity polymers into high-performance polyarylate materials with direct utilization of capping agent impurities. Nature Communications 16, 2482, doi:10.1038/s41467-025-57821-7 (2025).

32. Tanunchai, B., Nonthijun, P., Schädler, M., Disayathanoowat, T. & Noll, M. The enrichment of nitrogen-fixing bacteria on biodegradable plastics during the early stage of degradation under agricultural soil conditions and changing climate. Journal of Hazardous Materials Advances 20, 100793, 10.1016/j.hazadv.2025.100793 (2025).

33. Chen, D. et al. Upcycling of expanded polystyrene waste: Amination as adsorbent to recover Eriochrome Black T and Congo red. Separation and Purification Technology 289, 120669, 10.1016/j.seppur.2022.120669 (2022).

34. Liu, Y. et al. Atmospheric warming contributions from airborne microplastics and nanoplastics. Nature Climate Change, doi:10.1038/s41558-026-02620-1 (2026).

35. Choi, S., Lee, S., Kim, M.-K., Yu, E.-S. & Ryu, Y.-S. Challenges and Recent Analytical Advances in Micro/Nanoplastic Detection. Analytical Chemistry 96, 8846–8854, doi:10.1021/acs.analchem.3c05948 (2024).

36. Vico, C. & Chua, S. L. Optical, Chemical, and Biological Detection Methods of Microplastics and Nanoplastics. ACS Measurement Science Au, doi:10.1021/acsmeasuresciau.6c00028 (2026).

37. Mandemaker, L. D. & Meirer, F. Spectro-microscopic techniques for studying nanoplastics in the environment and in organisms. Angewandte Chemie International Edition 62, e202210494 (2023).

38. Materić, D. Nanoplastics measurements must have appropriate blanks. Proc Natl Acad Sci U S A 121, e2411099121, doi:10.1073/pnas.2411099121 (2024).

39. Sun, Y., Jiao, X., Zhang, N., Yan, B. & Fan, D. Correspondence on “Assessing the Release of Microplastics and Nanoplastics from Plastic Containers and Reusable Food Pouches: Implications for Human Health”. Environmental Science & Technology 58, 9013–9014, doi:10.1021/acs.est.4c02467 (2024).

40. Liu, Z. et al. Effects of nanoplastics at predicted environmental concentration on Daphnia pulex after exposure through multiple generations. Environ Pollut 256, 113506, doi:10.1016/j.envpol.2019.113506 (2020).

41. Tan, Y., Zhu, X., Wu, D., Song, E. & Song, Y. Compromised Autophagic Effect of Polystyrene Nanoplastics Mediated by Protein Corona Was Recovered after Lysosomal Degradation of Corona. Environ Sci Technol 54, 11485–11493, doi:10.1021/acs.est.0c04097 (2020).

42. Nie, J. H. et al. Polystyrene nanoplastics exposure caused defective neural tube morphogenesis through caveolae-mediated endocytosis and faulty apoptosis. Nanotoxicology 15, 885–904, doi:10.1080/17435390.2021.1930228 (2021).

43. Wei, W. et al. Anionic nanoplastic exposure induces endothelial leakiness. Nat Commun 13, 4757, doi:10.1038/s41467-022-32532-5 (2022).

44. Hsu, W. H. et al. Polystyrene nanoplastics disrupt the intestinal microenvironment by altering bacteria-host interactions through extracellular vesicle-delivered microRNAs. Nat Commun 16, 5026, doi:10.1038/s41467-025-59884-y (2025).

45. Cheng, S. et al. The effects of size and surface functionalization of polystyrene nanoplastics on stratum corneum model membranes: An experimental and computational study. Journal of Colloid and Interface Science 638, 778–787, 10.1016/j.jcis.2023.02.008 (2023).

46. Pitt, J. A. et al. Uptake, tissue distribution, and toxicity of polystyrene nanoparticles in developing zebrafish (*Danio rerio*). Aquat Toxicol 194, 185–194, doi:10.1016/j.aquatox.2017.11.017 (2018).

47. Pinsino, A. et al. Amino-modified polystyrene nanoparticles affect signalling pathways of the sea urchin (Paracentrotus lividus) embryos. Nanotoxicology 11, 201–209, doi:10.1080/17435390.2017.1279360 (2017).

48. Greven, A. C. et al. Polycarbonate and polystyrene nanoplastic particles act as stressors to the innate immune system of fathead minnow (Pimephales promelas). Environ Toxicol Chem 35, 3093–3100, doi:10.1002/etc.3501 (2016).

49. Mattsson, K. et al. Brain damage and behavioural disorders in fish induced by plastic nanoparticles delivered through the food chain. Sci Rep 7, 11452, doi:10.1038/s41598-017-10813-0 (2017).

50. Canesi, L. et al. Evidence for immunomodulation and apoptotic processes induced by cationic polystyrene nanoparticles in the hemocytes of the marine bivalve Mytilus. Mar Environ Res 111, 34–40, doi:10.1016/j.marenvres.2015.06.008 (2015).

51. Møller, P. & Roursgaard, M. Exposure to nanoplastic particles and DNA damage in mammalian cells. Mutation Research - Reviews in Mutation Research 792, 108468, 10.1016/j.mrrev.2023.108468 (2023).

52. Gopinath, P. M. et al. Assessment on interactive prospectives of nanoplastics with plasma proteins and the toxicological impacts of virgin, coronated and environmentally released-nanoplastics. Scientific Reports 9, 8860, doi:10.1038/s41598-019-45139-6 (2019).

53. Halimu, G. et al. Toxic effects of nanoplastics with different sizes and surface charges on epithelial-to-mesenchymal transition in A549 cells and the potential toxicological mechanism. J Hazard Mater 430, 128485, doi:10.1016/j.jhazmat.2022.128485 (2022).

54. Zinchenko, A. A., Sakaue, T., Araki, S., Yoshikawa, K. & Baigl, D. Single-Chain Compaction of Long Duplex DNA by Cationic Nanoparticles: Modes of Interaction and Comparison with Chromatin. The Journal of Physical Chemistry B 111, 3019–3031, doi:10.1021/jp067926z (2007).

55. Laemmli, U. K. Characterization of DNA condensates induced by poly(ethylene oxide) and polylysine. Proc Natl Acad Sci U S A 72, 4288–4292, doi:10.1073/pnas.72.11.4288 (1975).

56. Kabanov, V. A. et al. Interpolyelectrolyte Complexes Formed by DNA and Astramol Poly(propylene imine) Dendrimers. Macromolecules 33, 9587–9593, doi:10.1021/ma000674u (2000).

57. Anees, F., Montoya, D. A., Pisetsky, D. S. & Payne, C. K. DNA corona on nanoparticles leads to an enhanced immunostimulatory effect with implications for autoimmune diseases. Proc Natl Acad Sci U S A 121, e2319634121, doi:10.1073/pnas.2319634121 (2024).

58. Zeiger, E. The test that changed the world: The Ames test and the regulation of chemicals. Mutat Res Genet Toxicol Environ Mutagen 841, 43–48, doi:10.1016/j.mrgentox.2019.05.007 (2019).

59. Ames, B. N., Lee, F. D. & Durston, W. E. An improved bacterial test system for the detection and classification of mutagens and carcinogens. Proc Natl Acad Sci U S A 70, 782–786, doi:10.1073/pnas.70.3.782 (1973).

60. Duardo, R. C., Guerra, F., Pepe, S. & Capranico, G. Non-B DNA structures as a booster of genome instability. Biochimie 214, 176–192, 10.1016/j.biochi.2023.07.002 (2023).

61. Krzyżewska-Dudek, E. et al. The Influence of Lipopolysaccharide O-Antigen Chain Length on Biofilm Formation Capacity and Outer Membrane Proteome Shape of Salmonella Enteritidis. Environmental Microbiology Reports 17, e70211, 10.1111/1758-2229.70211 (2025).

62. Harley-Nyang, D., Memon, F. A., Osorio Baquero, A. & Galloway, T. Variation in microplastic concentration, characteristics and distribution in sewage sludge & biosolids around the world. Science of The Total Environment 891, 164068, 10.1016/j.scitotenv.2023.164068 (2023).

63. Muñiz, R. & Rahman, M. S. Microplastics in coastal and marine environments: A critical issue of plastic pollution on marine organisms, seafood contaminations, and human health implications. Journal of Hazardous Materials Advances 18, 100663, 10.1016/j.hazadv.2025.100663 (2025).

64. 64. Parmentier, Y., Bossant, M. J., Bertrand, M. & Walther, B. in Comprehensive Medicinal Chemistry II (eds John B. Taylor & David J. Triggle) 231-257 (Elsevier, 2007).

65. Moscatiello, G. Y. et al. The surface charge both influences the penetration and safety of polystyrene nanoparticles despite the protein corona formation. Environmental Science: Nano 12, 2857–2870, doi:10.1039/D4EN00962B (2025).

66. Kim, S. Y., Kim, Y. J., Lee, S.-W. & Lee, E.-H. Interactions between bacteria and nano (micro)-sized polystyrene particles by bacterial responses and microscopy. Chemosphere 306, 135584, 10.1016/j.chemosphere.2022.135584 (2022).

67. Dai, S. et al. Distinct lipid membrane interaction and uptake of differentially charged nanoplastics in bacteria. J Nanobiotechnology 20, 191, doi:10.1186/s12951-022-01321-z (2022).

68. Mafla-Endara, P. M. et al. Exposure to polystyrene nanoplastics reduces bacterial and fungal biomass in microfabricated soil models. Science of The Total Environment 904, 166503, 10.1016/j.scitotenv.2023.166503 (2023).

69. Strahl, H. & Hamoen, L. W. Membrane potential is important for bacterial cell division. Proceedings of the National Academy of Sciences 107, 12281–12286, doi:doi:10.1073/pnas.1005485107 (2010).

70. Wang, G. & Vasquez, K. M. Dynamic alternative DNA structures in biology and disease. Nat Rev Genet 24, 211–234, doi:10.1038/s41576-022-00539-9 (2023).

71. Arnott, S. & Selsing, E. The conformation of C-DNA. J Mol Biol 98, 265–269, doi:10.1016/s0022-2836(75)80115-x (1975).

72. Kypr, J., Kejnovská, I., Renciuk, D. & Vorlícková, M. Circular dichroism and conformational polymorphism of DNA. Nucleic Acids Res 37, 1713–1725, doi:10.1093/nar/gkp026 (2009).

73. Potaman, V. N., Alexeev, D. G., Skuratovskii, I., Rabinovich, A. Z. & Shlyakhtenko, L. S. Study of DNA films by the CD, X-ray and polarization microscopy techniques. Nucleic Acids Res 9, 55–64, doi:10.1093/nar/9.1.55 (1981).

74. Premilat, S. & Albiser, G. A new D-DNA form of poly(dA-dT).poly(dA-dT): an A- DNA type structure with reversed Hoogsteen pairing. Eur Biophys J 30, 404–410, doi:10.1007/s002490100170 (2001).

75. Subramani, V. K. & Kim, K. K. Characterization of Z-DNA Using Circular Dichroism. Methods Mol Biol 2651, 33–51, doi:10.1007/978-1-0716-3084-6_2 (2023).

76. Zaichuk, T. & Marko, J. F. Single-molecule micromanipulation studies of methylated DNA. Biophys J 120, 2148–2155, doi:10.1016/j.bpj.2021.03.039 (2021).

77. Lang, M. C., Malfoy, B., Freund, A. M., Daune, M. & Leng, M. Visualization of Z sequences in form V of pBR322 by immuno-electron microscopy. Embo j 1, 1149–1153, doi:10.1002/j.1460-2075.1982.tb00005.x (1982).

78. Fish, S. R., Chen, C. Y., Thomas, G. J., Jr. & Hanlon, S. Conformational characteristics of deoxyribonucleic acid-butylamine complexes with C-type circular dichroism spectra. 2. A Raman spectroscopic study. Biochemistry 22, 4751-4756, doi:10.1021/bi00289a021 (1983).

79. Buske, F. A., Mattick, J. S. & Bailey, T. L. Potential in vivo roles of nucleic acid triple-helices. RNA Biol 8, 427–439, doi:10.4161/rna.8.3.14999 (2011).

80. Kim, J. H., Kim, H. & Ko, K. S. Impact of DNA methyltransferases on bacterial fitness and genome stability in *Escherichia coli*. Journal of Global Antimicrobial Resistance 46, 203–208, 10.1016/j.jgar.2025.12.009 (2026).

81. Bhagwat, A. S. & Roberts, R. J. Genetic analysis of the 5-azacytidine sensitivity of *Escherichia coli* K-12. J Bacteriol 169, 1537–1546, doi:10.1128/jb.169.4.1537-1546.1987 (1987).

82. Jin, D. J. & Gross, C. A. Mapping and sequencing of mutations in the *Escherichia coli rpoB* gene that lead to rifampicin resistance. J Mol Biol 202, 45–58, doi:10.1016/0022-2836(88)90517-7 (1988).

83. Banyay, M., Sarkar, M. & Gräslund, A. A library of IR bands of nucleic acids in solution. Biophysical Chemistry 104, 477–488, 10.1016/S0301-4622(03)00035-8 (2003).

84. Anifowoshe, Abass T. et al. Genotoxicity and Genomic Instability Induced by Micro- and Nanoplastics: A Comprehensive Multi-Taxa Mechanistic Review. Journal of Applied Toxicology 46, 1750–1778, 10.1002/jat.70122 (2026).

85. Sadique, S. A. et al. Impact of microplastics and nanoplastics on human Health: Emerging evidence and future directions. Emerging Contaminants 11, 100545, 10.1016/j.emcon.2025.100545 (2025).

86. Paget, V. et al. Specific uptake and genotoxicity induced by polystyrene nanobeads with distinct surface chemistry on human lung epithelial cells and macrophages. PLoS One 10, e0123297, doi:10.1371/journal.pone.0123297 (2015).

87. Gelova, S. P. & Chan, K. Mutagenesis induced by protonation of single-stranded DNA is linked to glycolytic sugar metabolism. Mutat Res 826, 111814, doi:10.1016/j.mrfmmm.2023.111814 (2023).

88. Sowers, L. C., David Sedwick, W. & Shaw, B. R. Hydrolysis of N3-methyl-2′- deoxycytidine: Model compound for reactivity of protonated cytosine residues in DNA. Mutation Research - Fundamental and Molecular Mechanisms of Mutagenesis 215, 131–138, 10.1016/0027-5107(89)90225-X (1989).

89. Cárdenas, M., Schillén, K., Nylander, T., Jansson, J. & Lindman, B. DNA Compaction by cationic surfactant in solution and at polystyrene particle solution interfaces: a dynamic light scattering study. Physical Chemistry Chemical Physics 6, 1603–1607, doi:10.1039/B310798A (2004).

90. Mortelmans, K. & Zeiger, E. The Ames *Salmonella*/microsome mutagenicity assay. Mutat Res 455, 29–60, doi:10.1016/s0027-5107(00)00064-6 (2000).

91. Baba, T. et al. Construction of *Escherichia coli* K-12 in-frame, single-gene knockout mutants: the Keio collection. Mol Syst Biol 2, 2006 0008, doi:10.1038/msb4100050 (2006).

92. Bale, A., d’Alarcao, M. & Marinus, M. G. Characterization of DNA adenine methylation mutants of *Escherichia coli* K-12. Mutation Research/Fundamental and Molecular Mechanisms of Mutagenesis 59, 157–165, 10.1016/0027-5107(79)90153-2 (1979).

93. Bolivar, F. et al. Construction and characterization of new cloning vehicles. II. A multipurpose cloning system. 1977. Biotechnology 24, 153-171 (1992).

94. Kang, M. & Kim, D. Crystallization of Z-DNA in Complex with Chemical and Z-DNA Binding Z-Alpha Protein. Methods Mol Biol 2651, 59–67, doi:10.1007/978-1-0716-3084-6_4 (2023).

