## Supplementary material for "Nanoplastic-mediated non-B DNA mutagenicity": Suppl Methods

**Supplementary Information**

**Materials and Methods:**

**Bacterial strains and growth conditions**. Description to all strains, plasmids, and oligos used in this work are listed in **Table 1**. Bacteria were cultured aerobically at 37°C in plain lysogeny broth (LB) or agar (LA) in glass-Erlenmeyer flasks. For the Ames-test, the top agar and minimal glucose medium were prepared according to Mortelmans and Zeiger (2000)[90]. Incubation of bacteria with 20μg/ml decitabine (5-aza-2′-deoxycytidine, HY-A0004R, MedChemExpress) and 50μM CCCP (carbonyl cyanide m-chlorophenylhydrazone; C2759, Merck) was performed aerobically at 37°C for 30min. In short, 3ml of overnight bacterial culture was spun down and washed 3 times with PBS (phosphate-buffered saline), the pellet was resuspended in 3ml LB with CCCP or decitabine. After incubation, the bacterial samples were washed again 3 times with PBS. For rescue experiments, 2% of MUTAZYME™ 10% S9 Mix (11-404L, TrinovaBiochem) and 0.1mg/ml NAC (N-acetylcyteine; A0150000, Merck) were included in the preincubation reactions in the Ames test. Bacteria that underwent stress-induced mutagenesis were plated on LA 100μg/ml rifampicin (R3501, Merck) for up to 72 hours.

**Characterization of nanoplastics.** Suspensions of 1% solid content polystyrene (PS), amine functional polystyrene (PS-$NH_2$), carboxyl functional polystyrene (PS-COOH), as well as polymethyl methacrylate (PMMA), amine functional polymethyl methacrylate (PMMA-$NH_2$), carboxyl functional polymethyl methacrylate (PMMA-COOH) microspheres with a 25nm diameter were purchased from Lab 261 (Palo Alto, CA, USA) with catalog numbers PST25, PST25A, PST25C, PMMA25, PMMA25A and PMMA25C, respectively, as well as PS-$NH_2$ with a diameter of 300 and 1000nm (catalog numbers PST300A and PST1000A, respectively). To remove antibacterial additives $NaN_3$ (2mM) and Tween-20 (0.1%), 3ml of each suspension was subjected to overnight dialysis in a Slide-A-Lyzer 3.5k cassette (Thermo Scientific, 66330) with a 3500-Da cut-off. To assess the quality of the nanoparticles after dialysis, each suspension was visualized via transmission electron microscopy (TEM).

**Mutagenic experiments with bacteria with Ames test.** The experiments were done based on the Mortelmans and Zeiger (2000)[1] protocol with autoclaving being replaced with sterile filtration with strains listed in **Table1**. In short, single colonies of *Salmonella*

TA1535 and TA1538 were cultured overnight at 37°C in replicates of three and five. Then, the preincubation assay was performed as follows: 15µl of 1% plastic solution and 35µl of PBS were added in each of the test tubes, while the control tubes contained only 50µl of PBS. Then, 100µl of washed $10^8$ bacteria were added into the designated tube. The samples were incubated for 1h aerobically at 37°C. The top agar was melted and aliquoted into 15-ml tubes and kept molten at 48°C. After incubation, the samples were mixed with the top agar, poured upon the designated glucose minimal agar and after solidification, the plates were incubated at 37°C for 48h. The revertants were counted compared to the ones on the control plates, as well as with the numbers of spontaneous revertants.

To assess the effect of nanoplastics on stress-induced mutagenesis in epigenetically silenced bacteria, we performed a rifampicin mutagenicity assay. This is a linear assay that allows the monitoring of neutral mutations since almost all the mutations responsible for rifampicin resistance are in the *rpoB* gene[2]. In short, 3ml of overnight bacterial culture were pelleted (3000xg, 10min), washed 3 times with PBS. Tubes containing 35µl of PBS and 15µl of 1% nanoplastic solution or PBS for controls were prepared and 100µl of washed $10^8$ bacteria were added into the designated tubes. The samples were incubated for 1h and overnight aerobically at 37°C, then plated on LA with and without rifampicin for up to 72 hours. The mutation frequency is displayed as a ratio between the resistant colonies and the total bacterial number.

**Electron microscopy (EM) of bacterium-nanoplastic interactions.** To visualize the bacterium-nanoplastic interactions, bacteria incubated with 0.1% NPs in 1XPBS for 1h at 37°C, were spun down and washed three times with cacodylate buffer (0.05M KCl, 0.005M $MgCl_2$, 0.067M cacodylate) at 6000xg for 15min. Then the pellets were resuspended and fixed overnight in 2.5% glutaraldehyde at 4°C. After fixation, bacteria were washed again with cacodylate buffer and embedded in 1% low-gradient agarose (prepared in cacodylate buffer). After postfixation in osmiumtetroxide and dehydration, bacterial pellets were embedded in resin (AGAR 100 Resin, Agar Scientific). Ultrathin sections (80nm) were contrasted with uranyl acetate/lead citrate and studied with a Zeiss EM 900 electron microscope.

**Electron microscopy of plasmid-nanoplastic complexes.** To visualize the V-form of pBR322, we performed immunoelectron microscopy of plasmid- PS-$NH_2$ complexes following the method used by Lang et al.[3]. In short, 1µg of pBR322 was exposed to 0.1% PS-$NH_2$ for 1h at 37°C in 100mM Tris-HCl buffer (pH 7.5). Then, together with the untreated control, the plasmid-NP complex was incubated with the primary anti-Z-DNA antibody (1:1000 dilution) for 1h at 25°C. The anti-Z-DNA antibody (Ab00783-1.29, Nordic Biosite) was raised against brominated poly(dG-dC)-poly(dG-dC) sequence in C57BL/6 mice. After incubation, the antibody-DNA complexes were gel filtrated via Sepharose ™ 4B (4B200, Merck) column, and incubated with immunogold labelled secondary anti-mouse antibody (G7652, Merck) in a 1:50 ratio for 2h at 25°C. The samples subjected to TEM were either gel-filtrated or not after incubation with the secondary antibody. Plasmid DNA in spreading solution (0.002% benzyl dimethyl alkyl ammonium chloride and 0.5M ammonium acetate) was adsorbed onto Formvar coated copper grids by placing them on a drop of 7µl plasmid DNA mixture for 3min. The grids were then washed briefly by touching them to the surface of three drops of distilled water. The washing step was followed by a pre-hydration step by touching grids to the surface of a drop of 50% methanol. The nucleic acid was stained by placing the grids onto a drop of 2% uranyl acetate in 70% methanol for 30sec. To rinse, grids were briefly placed on a drop of 70% methanol. After drying, grids are immediately ready for TEM studies with a Zeiss EM 900 electron microscope.

**Whole genome sequencing (WGS) of the Ames *Salmonella* strains.** Whole genomes were extracted from 10 spontaneous revertants of TA1535, 10 NP-induced revertants of TA1535, 10 spontaneous revertants of TA1538 and10 NP-induced revertants of TA1538. In short, bacteria from 10ml overnight culture were harvested and spun down at 8000xg at 4°C for 15min. The pellets were washed twice with normal saline and spun down at 8000xg for 15min. Then, they were resuspended in 0.5ml of 10mM Tris-HCl (pH 8) and 2.5mg/ml of lysozyme and incubated at 37°C for 1h. After incubation,1ml of lysis buffer (50 mM Tris, 100 mM EDTA, 1% SDS, pH 8) and 1mg/ml of proteinase-K were added to the samples, and they were incubated at 50°C for 1h in a water bath. The lysis was followed by the addition of 1ml of phenol:chloroform:isoamyl alcohol (25:24:1). Samples were mixed for 2-3 minutes and centrifuged at 10000xg at 4°C for 15min. The aqueous phase was transferred in a fresh sterile tube, and 50µl of 3M sodium acetate was added, together with twice the volume of 95% ice-cold ethanol. The precipitation was carried out for 1h at -20°C. After settling down, samples were centrifuged at 10000xg at 4°C for 15min. The supernatants were decanted, and the pellets washed with 70% ethanol and centrifuged again. After drying, the pellets were dissolved in water, and the quality and quantity of the obtained DNA measured with NanoDrop Lite Plus micro-UV spectrophotometer (ThermoFischer

Scientific). 100ng of each sample was subjected to WGS. PCR-free sequencing libraries were prepared from genomic DNA using the xGen EZ UNI library preparation kit (Integrated DNA Technologies) according to the manufacturer's instructions. During adapter ligation, KAPA Universal Adapters (Roche) were used for sample indexing and multiplexing. Libraries were sequenced on the Element AVITI system (Element Biosciences) in paired-end mode (2X150 bp). Sequencing was performed to a target depth of at least 100× genome coverage per sample. The genomes of the parental as well as the NP-generated and spontaneous mutants/revertants are deposited in the public sequence database GenBank under the BioProject ID: PRJNA1515829 (samples TA1535C1-TA1535C10) and PRJNA1519092 (samples TA1535PS1-TA1535PS10; TA1538C1- TA1538C10; TA1538PS1- TA1538PS10; TA1535 and TA1538).

**Variant calling analysis.** Raw sequencing reads were quality-checked using FastQC, pre-processed with Cutadapt v4.3,1 and aligned to the *Salmonella enterica* subsp. *enterica* serovar Typhimurium strain LT2 reference genome (GenBank assembly ASM694v2; accession GCF_000006945.2) using BWA-MEM v0.7.17.2 Duplicate reads were identified and flagged using Picard MarkDuplicates v2.27.1. To characterize the variant background of each parental strain relative to the reference, somatic variant calling was performed in tumor-only mode by employing an ensemble of three callers: Strelka v2.9.10,3 VarDict v.1.8.3,4 and VarScan v2.4.4.5 This approach was favored over germline calling to avoid imposing fixed-allele-frequency priors, which may not be applicable to laboratory stock populations that have experienced drift or incomplete clonal fixation. The resulting variant records were employed as the parental background reference for subsequent analyses. Variants in experimental samples were identified using a paired somatic design, designating each experimental sample as the tumor counterpart and its corresponding parental strain as the matched normal. Paired variant calling was conducted employing the same three-caller ensemble. Therefore, variants inherent to the parental background are systematically modelled and excluded from the calls, thereby retaining only mutations that are independently acquired in each experimental lineage.

**Unique mutation and motif discovery.** We selected unique mutations by merging variant calls across biological replicates within each experimental group. Genomic sequences of 100 nucleotides upstream (5′) and downstream (3′) of each mutation site were extracted from the reference genome. GC content was calculated for each upstream and downstream sequence. Differences between spontaneous and nanoparticle-induced revertants were evaluated using two-sided Mann–Whitney U tests with Benjamini–Hochberg correction for multiple testing.

*De novo* sequence motifs enriched in mutation-associated regions were identified using the MEME Suite (v5.5.9) (PMID: 25953851). Motif discovery was performed separately for upstream and downstream flanking regions using MEME under the zero-or-one-occurrence-per-sequence (ZOOPS) model with motif widths ranging from 6 to 20 nucleotides. A matched genomic background was generated by randomly sampling genomic regions of identical length from the same chromosome. Similar numbers and lengths of tested sequences and randomly selected background sequences were used to control chromosome-specific compositional biases. Position weight matrices (PWM) from MEME were then used as input for FIMO to scan both mutation-associated and matched background sequences. Motif enrichment was assessed by comparing the frequency of motif occurrences between tested and background datasets using the significance thresholds implemented in FIMO. Consensus motifs were visualized as sequence logos generated from the corresponding PWM.

**Positional association of mutations with predicted non-B-DNA motifs.** For each substitution, the absolute genomic distance to the nearest predicted non-B-DNA motif was calculated from the corresponding genomic coordinates. The distribution of observed mutation-to-motif distances was compared with a null distribution generated from an equal number of randomly sampled positions across the reference genome (AE006468.2. Empirical cumulative distribution functions (ECDFs) were used to visualize differences in motif proximity between observed mutation sites and random genomic positions. Differences between the observed and random distance distributions were done by using a one-sided Mann–Whitney U test to observe if mutation sites occurred closer to predicted non-B-DNA motifs are due to expected by chance. Motif-centered mutation-density profiles were generated by centering the predicted non-B-DNA motifs at position zero, and the distribution of mutations within flanking intervals around each motif were calculated. The non-B-DNA sequences were predicted with Z-hunter, with the following parameters: minimal sequence size: 8; minimal score (%): 50%; Score GC: 50; Score GT/AC: 3 with estimated frequency per 1000 kb (https://bioinformatics.ibp.cz/#/).

**Spectral measurements- Circular dichroism (CD).** CD was measured within the range of 330 - 220 nm with J-1700 CD spectrometer. To generate CG and BZ DNA heteroduplexes, the complementary strands of the DNA were annealed as follows: 95°C, 90°C, 80°C, 70°C, 60°C, 50°C, 40°C, 30°C, 20°C, and10°C for 10min each step. Prior to measurements, 20ng/µl template (or 10ng/µl chromosomal DNA) was either mixed with 0.1% NP solution and transferred to 10-mm quartz cuvette or directly used as a control. Each measurement was done three times in technical replicates, and each spectrum in the figure represents the median of these measurements. For samples from biological experiments, in addition to the technical replicates, biological replicates were included.

**FT-IR (Fourier transform infrared spectroscopy) microspectroscopy.** FT-IR of DNA-NP complexes was measured with Perkin Elmer Frontier IR microscope. DNA-NP complexes were extracted from $10^9$ bacteria incubated for 1h with 0.1% NPs in PBS. The bacteria were pelleted (13000xg, 15 min) and washed 3 times with PBS and once with 10mM Tris-HCl (pH 8.0). The washed bacteria were subjected to cell fractionation. In short, bacteria were incubated for 10min twice and washed in 20mM Tris-HCl, 20% sucrose and 0.1mM EDTA. The pellet was then incubated in water on ice for 10min and spun down. The supernatant contained the bacterial periplasmic fraction. The pellet (bacterial protoplasts) was lysed with 150µl lysis buffer (50mM Tris-HCl, 1% sodium dodecyl sulfate (SDS), 100mM EDTA) at 50°C for 1h and DNA precipitated with ice-cold ethanol. The pellets (DNA-NP complexes) were subjected to IR spectroscopy and thermogravimetric experiments. For the DNA release assay, the pellets were incubated in 150µl water and heated up for 1h at 50°C.

Both attenuated reflectance (ATR) infrared (IR) spectroscopy and micro-Fourier transform (FT) IR spectroscopy have been used. A Perkin Elmer Spotlight IR microscope was used for micro FTIR measurements on the extracellular environment, of which 10µl was dropcasted on a 13mm diameter round calcium fluoride window (Spectran), which was subsequently heated on a heating plate at 80°C until dry. They were measured in transmission mode (2 $cm^{-1}$ resolution, 16 averages) with a background recording on fresh calcium fluoride. Before every scan, the background was validated by doing a measurement outside the dried sample drop. The samples were measured on the dried outer ring to yield proper intensity. For the IR measurements shown in Figure 3 a Perkin Elmer 2000 with ATR-module has been used with 2 $cm^{-1}$ resolution and 32 averages.

**Statistical analysis.** All the experiments were repeated in biological replicates (Ames test, stress-induced mutagenesis, and EM) and 10 biological replicates for the WGS.

The statistical analysis of the experiments was performed with GraphPad Prism 10. Pairwise comparisons of samples were done using unpaired Welch´s t-tests assuming normal or lognormal distributions with multiple t-test with Welch corrections, one-way ANOVA with Tukey's correction and two-sided Mann–Whitney U tests with Benjamini–Hochberg correction for multiple testing. Significance was determined by a P-value < 0.05, as follows: P > 0.05 (ns, i.e., not significant); P ≤ 0.05 (*); P ≤ 0.01 (**); P ≤ 0.001 (***).

**Table 1. Bacterial strains, plasmids and oligos used in this work.**

| **Strains, plasmids and oligonucleotide primers used** | **Description/Relevant characteristics** | **Reference** |
|---|---|---|
| ***Salmonella enterica*** | | |
| TA1535 | *rfa*$^-$, *ΔuvrB, hisG46* | 4 |
| TA1538 | *rfa*$^-$, *ΔuvrB, hisD3052* | 4 |
| ***Escherichia coli*** | | |
| BW25113 | F$^-$, Δ(*araD-araB*)567, *ΔlacZ*4787(::*rrnB*-3), λ$^-$, *rph-1*, *Δ(rhaD-rhaB)568*, *hsdR514* | 5 |
| JW1944-2 | F$^-$, Δ(*araD-araB*)567, *ΔlacZ*4787(::*rrnB*-3), λ$^-$, *Δdcm-735::kan*, *rph-1*, *Δ(rhaD-rhaB)568*, *hsdR514* | 5 |
| GM99 | F-, *dam4, mal 354, tsx354* | 6 |
| GM2163 | F–, *ara-14*, *leuB6*, *fhuA31*, *lacY1*, *tsx78*,*glnV44*, *galK2*, *galT22*, *mcrA*, *dcm-6*, *hisG4*, *rfbD1*, *rpsL136*, *dam13*::Tn9, *xylA5*, *mtl–1*, *thi-1*, *mcrB1*, *hsdR2* | 6 |
| **Plasmids** | **Description/Relevant characteristics** | **Reference** |
| pBR322 | Cb$^R$, Tc$^R$, *rep* ori (pMB1) | 7 |
| **Oligonucleotides** | **Sequence (5'->3'):** | |
| BZFw | CGCGCGCGCGCGATAAACCACTCGG | 8 |
| BZRv | CCGAGTGGTTTATCGCGCGCGCGCG | 8 |
| CGFw | CGCGCGCGCGCG | 8 |
| CGRv | CGCGCGCGCGCG | 8 |
| t(cg)3 | TCGCGCG | 9 |
| (cg)3a | CGCGCGA | 9Rif |
| C(Me)GFw | G[5MeC]G[5MeC]G[5MedC]G[5MedC]G[5MedC]G | This study |
| C(Me)GRv | [5MedC]G[5MedC]G[5MedC]G[5MedC]G[5MedC]G[5MedC] | This study |
| ACFw | ACACACACACAC | This study |
| ACRv | GTGTGTGTGTGT | This study |
| ATFw | ATATATATATAT | This study |

| | | |
|---|---|---|
| ATRv | ATATATATATAT | This study |
| ZDNAFW | CGCGCGCGCGCGCGCGCGCGCGCGCGC GCGCGCGCGCGCGCGCGCGCGCG | This study |
| ZDNARV | CGCGCGCGCGCGCGCGCGCGCGCGCGC GCGCGCGCGCGCGCGCGCGCGCG | This study |
| polydCTFw | CTCTCTCTCTCT | This study |
| polydCTRv | AGAGAGAGAGAG | This study |
| polydTTCFw | TTCTTCTTCTTC | This study |
| polydTTCRv | GAAGAAGAAGAA | This study |
| polydTCCFw | TCCTCCTCCTCC | This study |
| polydTCCRv | GGAGGAGGAGGA | This study |

**References:**


1. Mortelmans, K. & Zeiger, E. The Ames *Salmonella*/microsome mutagenicity assay. Mutat Res 455, 29-60, doi:10.1016/s0027-5107(00)00064-6 (2000).

2. Jin, D. J. & Gross, C. A. Mapping and sequencing of mutations in the *Escherichia coli rpoB* gene that lead to rifampicin resistance. J Mol Biol 202, 45-58, doi:10.1016/0022-2836(88)90517-7 (1988).

3. Lang, M. C., Malfoy, B., Freund, A. M., Daune, M. & Leng, M. Visualization of Z sequences in form V of pBR322 by immuno-electron microscopy. Embo j 1, 1149-1153, doi:10.1002/j.1460-2075.1982.tb00005.x (1982).

4. Ames, B. N., Lee, F. D. & Durston, W. E. An improved bacterial test system for the detection and classification of mutagens and carcinogens. Proc Natl Acad Sci U S A 70, 782-786, doi:10.1073/pnas.70.3.782 (1973).

5. Baba, T. et al. Construction of *Escherichia coli* K-12 in-frame, single-gene knockout mutants: the Keio collection. Mol Syst Biol 2, 2006 0008, doi:10.1038/msb4100050 (2006).

6. Bale, A., d'Alarcao, M. & Marinus, M. G. Characterization of DNA adenine methylation mutants of *Escherichia coli* K-12. Mutation Research/Fundamental and Molecular Mechanisms of Mutagenesis 59, 157-165, doi:https://doi.org/10.1016/0027-5107(79)90153-2 (1979).

7. Bolivar, F. et al. Construction and characterization of new cloning vehicles. II. A multipurpose cloning system. 1977. Biotechnology 24, 153-171 (1992).

8. Subramani, V. K. & Kim, K. K. Characterization of Z-DNA Using Circular Dichroism. Methods Mol Biol 2651, 33-51, doi:10.1007/978-1-0716-3084-6_2 (2023).

9. Kang, M. & Kim, D. Crystallization of Z-DNA in Complex with Chemical and Z-DNA Binding Z-Alpha Protein. Methods Mol Biol 2651, 59-67, doi:10.1007/978-1-0716-3084-6_4 (2023).
